# Transsynaptic labeling and transcriptional control of zebrafish neural circuits

**DOI:** 10.1101/2023.04.03.535421

**Authors:** Cagney Coomer, Daria Naumova, Mustafa Talay, Bence Zolyomi, Nathaniel J. Snell, Altar Sorkaç, Jean-Michale Chanchu, Ji Cheng, Ivana Roman, Jennifer Li, Drew Robson, Gilad Barnea, Marnie E. Halpern

## Abstract

Deciphering the connectome, the ensemble of synaptic connections that underlie brain function, is a central goal of neuroscience research. Here, we report mapping of connections between presynaptic and postsynaptic partners in a living vertebrate nervous system, that of the zebrafish, through the successful adaptation of the *trans*-Tango genetic approach, first developed for anterograde transsynaptic tracing in *Drosophila*. Neural connections were visualized between synaptic partners in the larval retina and brain and followed over development. Results were corroborated by functional experiments in which optogenetic activation of retinal ganglion cells elicited responses in neurons of the optic tectum, as measured by *trans*-Tango-dependent expression of a genetically encoded calcium indicator.

Transsynaptic signaling through *trans*-Tango reveals predicted as well as previously undescribed synaptic connections in the zebrafish brain, providing a valuable *in vivo* tool to monitor and interrogate neural circuits over time.

## Introduction

Ever since the application of Golgi staining to the nervous system, neuroscientists have sought to understand the structural organization of the brain and its underlying neural connections (^1^). Labeling with dyes or fluorescent proteins, transsynaptic tracing through application of viruses and, more recently, reconstruction by serial electron microscopy (EM) have helped resolve aspects of the connectome in invertebrate and vertebrate models. Each approach has its benefits and limitations (^2^). For example, anterograde and retrograde viral tracers have been widely used to map neural connections in the mouse brain (^3–5^), but transduction of virus is variable in different cell types and inevitably causes death of infected neurons. The labor-intensive sample collection and computational complexity required for 3-D EM reconstruction, (^2, 6^), precludes surveying large brain regions or comparisons between samples.

Genetic methods for anterograde transsynaptic tracing permit precise labeling from identified presynaptic neuronal populations in multiple individuals and can be used to both identify synaptic partners and modify their activity. The genetically encoded synthetic transsynaptic labeling system, *trans*-Tango (^7^) has been successfully used to map a variety of neural circuits in *Drosophila* (^8–12^). The Tango system relies on a synthetic signaling pathway involving two fusion proteins that convert the activation of a receptor into reporter expression (^13^). To determine its effectiveness in a vertebrate central nervous system (CNS), we modified the components of *Drosophila trans*-Tango for anterograde transsynaptic labeling in zebrafish, a powerful vertebrate model for correlating whole brain imaging of neural activity with behavior.

Large numbers of zebrafish embryos can be readily injected at the 1-cell stage with plasmids bearing all *trans*-Tango components, and subsequent neuronal labeling assessed in the larval brain. Such transient assays enable a variety of constructs to be rapidly tested, compared, and optimized. The transparency of larval stages permits live confocal imaging of uniquely labeled pre and postsynaptic neurons throughout the brain and connections can be followed over time. Notably, neuronal morphologies are distinguished at the single-cell level by sparse labeling resulting from mosaic expression of injected DNA constructs. Many transgenic tools are also already available that can be used in conjunction with the *trans*-Tango reagents. As first described by Talay et al., (^7^), the *trans*-Tango approach relies on two binary transcriptional regulatory systems, Gal4/UAS from *Saccharomyces cerevisiae* and QF/QUAS from *Neurospora crassa* (Fig. 1a), that both function in zebrafish (^14, 15^). Adopting *trans*-Tango for zebrafish capitalizes on existing Gal4 lines to drive expression of a ligand derived from human glucagon (hGCG) in presynaptic neurons of interest. In this construct, hGCG is fused to the cytosolic and transmembrane domains of a zebrafish synaptic protein, either *neurexin 1a* or *1b*, and the extracellular domain of the *human intercellular cell adhesion molecule 1* (ICAM1). With these modifications, the ligand is tethered to the presynaptic membrane and extends into the synaptic cleft. The corresponding G protein-coupled glucagon receptor (hGCGR) is expressed on the surface of all neurons under control of a nearly pan neural promoter. The cytoplasmic tail of the receptor is fused to the QF transcription factor via a sequence containing the cleavage site for the N1a protease from the tobacco etch virus (TEV) (^13^). Upon binding of the ligand across the synapse, the activated G protein-coupled receptor recruits humanβ-arrestin2 that is fused to the N1a protease of the tobacco etch virus (hArr-TEV). Recruitment of hArr-TEV results in proteolytic cleavage of the TEV cleavage site (TEVcs) and release of the receptor bound QF transcription factor in postsynaptic neurons. QF can then translocate to the nucleus where it activates transcription of any gene downstream of the multicopy upstream activating sequence (QUAS). As we demonstrate, different reporters can thus be placed under UAS and QUAS control to label pre and postsynaptic neurons, respectively, with distinct fluorescent proteins. Additionally, target neurons can express any gene under QUAS control such as genetically encoded calcium indicators (i.e., GCaMP).

**Fig. 1:**
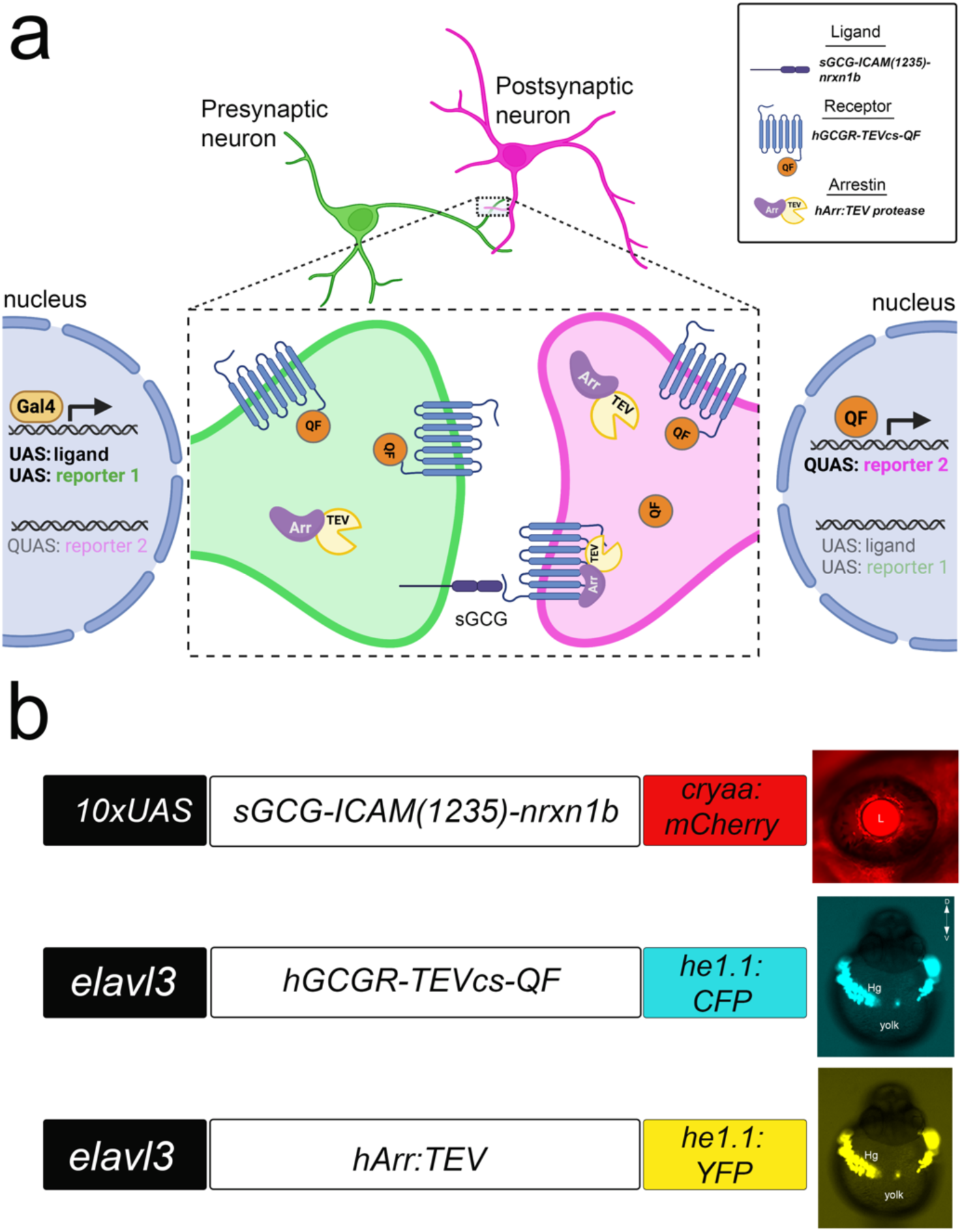
t*r*ans-Tango mediates transsynaptic signaling in zebrafish (a) Gal4 driver lines direct ligand expression in presynaptic neurons of interest. Ligand derived from human glucagon (sGCG) is tethered to the presynaptic membrane through the transmembrane domain of zebrafish Nrxn1b and extended into the synaptic cleft through the ICAM(1235) linker sequence (Talay et al., 2017). The receptor (hGCGR) is fused to the QF transcription factor through the protease cleavage site (cs) of the N1a protease from the tobacco etch virus (TEV) and is expressed by neurons throughout the CNS. Ligand binding to the receptor activates the pathway in postsynaptic neurons, resulting in recruitment of an Arrestin-TEV protease fusion protein and proteolytic cleavage of QF. QF can then translocate to the nucleus where it promotes transcription of genes downstream of the upstream activation sequence (QUAS) to which it binds. UAS and QUAS regulated reporters label pre and postsynaptic neurons, respectively, with green and red fluorescent proteins. (b) Constructs containing modified ligand, receptor and arrestin-TEV components were cloned into Tol2 transposition vectors. To confirm the presence of each *trans*-Tango component in injected embryos, secondary markers consisting of tissue specific promoters driving different FPs that label the lens of the eye (*cryaa*) or hatching gland cells (*he1.1*) were included in each Tol2 plasmid.

## Results

### Adaptation of *trans*-Tango components for zebrafish

To adapt the *trans*-Tango system for use in zebrafish, the ligand, receptor, and arrestin-TEV constructs were cloned into vectors containing the short flanking arms of the Tol2 transposon (^16^). The presence of each component was assessed by the addition of a secondary marker on the same plasmid, consisting of a tissue-specific promoter driving expression of a fluorescent protein (Fig. 1b). Six ligand constructs that differed in the length of the ICAM1 extracellular domain and in zebrafish *neurexin1a* or *1b* domains were placed under the control of multicopy UAS sequence (10xUAS) (^17^) in a Tol2 plasmid that also contained the *crystalline, alpha A* promoter driving mCherry (*cryaa:mCherry*). For transcription of the hGCGR-QF and hArr-TEV constructs, we used the promoter from the zebrafish *ELAV like neuron-specific RNA binding protein 3* (*elavl3*) gene that directs widespread expression in post-mitotic neurons throughout the larval zebrafish CNS (^18^). The receptor and arrestin-TEV constructs were cloned into Tol2 plasmids containing the promoter from the *hatching gland enzyme1.1* (*he1.1*) gene (^19^) upstream of sequences encoding cyan or yellow fluorescent protein, respectively. The *he1.1* promoter drives expression in hatching gland cells at 24 hours post fertilization (hpf); however, labeling is not detected after 72 hpf.

This permits imaging of neural connections in the larval brain unobscured by bright fluorescence from a secondary marker.

### Gal4-dependent anterograde labeling by *trans-*Tango

To assess the effectiveness of the individual ligand, receptor-QF, and arrestin-TEV constructs, we injected plasmids encoding each along with mRNA encoding Tol2 transposase into zebrafish embryos. The embryos were derived from matings between *pt1fa:Gal4-VP16* driver (^20^) and *QUAS:mApple-CAAX* reporter lines. The *pancreas-specific transcription factor 1a* (*ptf1a*) driver because it promotes transcription broadly in the hindbrain (^21–23^), as well as in retinal amacrine and horizontal cells and a subset of ganglion cells (^24, 25^), providing a substantial target area for testing the functionality of *trans* Tango. We determined which of six UAS:ligand constructs resulted in extensive labeling of putative postsynaptic neurons. Three constructs contained domains from the *neurexin 1a* gene and three from *neurexin 1b* (Supplementary Fig. 1) that either included the full-length extracellular domain from ICAM1, a truncated version, or lacked these sequences altogether (^7^). Criteria for determining the optimal ligand construct were mApple-CAAX reporter labeling that was robust, consistent between samples, and Gal4-dependent. We co-expressed the receptor and arrestin-TEV constructs with each ligand configuration and determined that 10x*UAS:sGCG-ICAM(1235)-nrxn1b* produced the most extensive mApple-CAAX labeling of target neurons with minimal Gal4-independent labeling (Supplementary Fig. 1). We, therefore, used this ligand construct for all subsequent experiments. In 89% of larvae that were screened positive for *pt1fa:Gal4-VP16* and all three *trans*-Tango components (n=1,042), mApple-CAAX labeled neurons were found in close proximity to GFP-labeled presynaptic neurons throughout the hindbrain (Fig. 2a,a’). In addition, we observed labeling of putative postsynaptic neurons in the optic tectum, presumably because the *pt1fa* promoter is active in a subset of retinal ganglion cells (^25^) whose axonal projections terminate in this midbrain region.

**Fig. 2:**
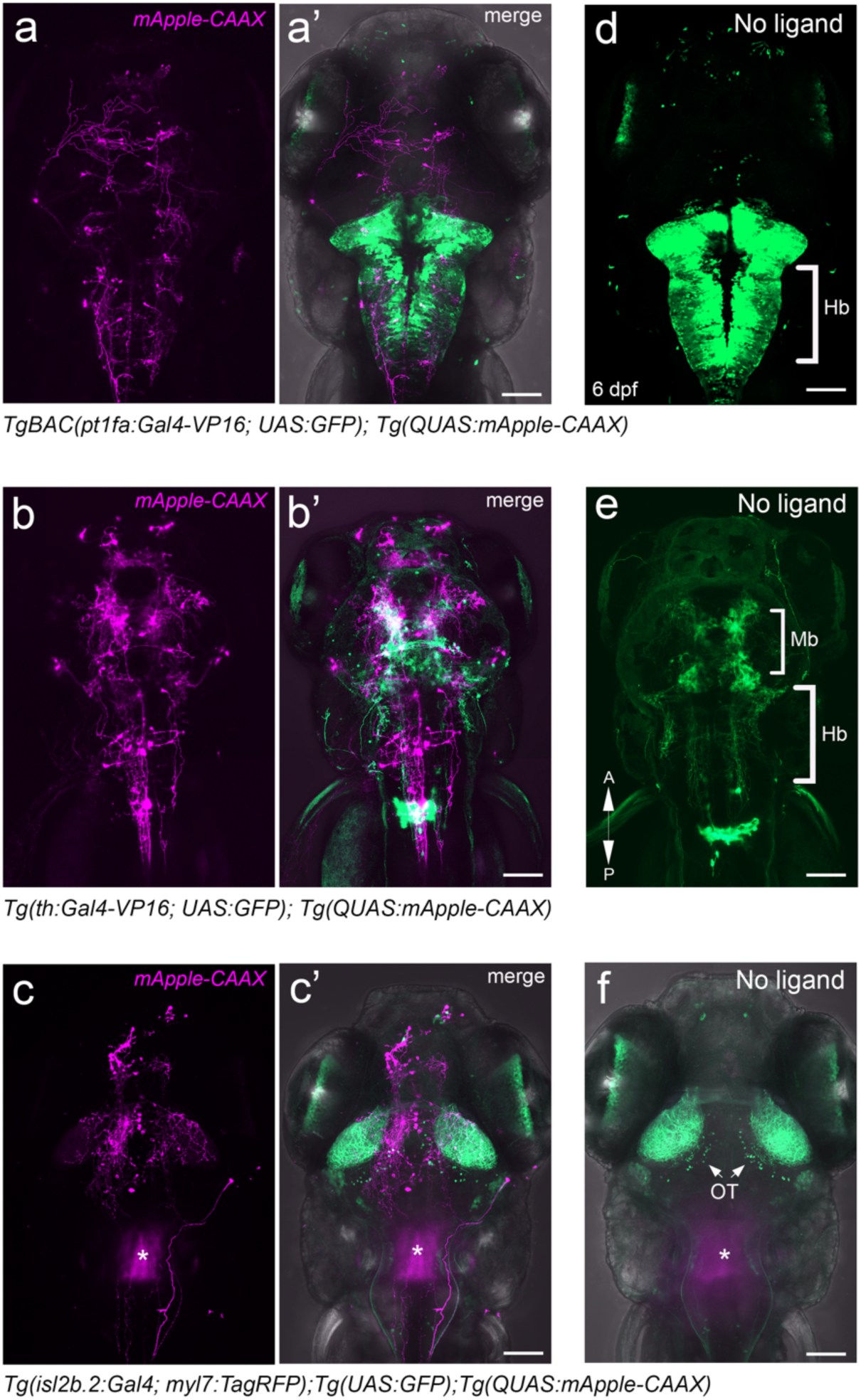
Gal4-dependent *trans*-Tango labeling of synaptic partners Progeny from Gal4 driver lines mated to *Tg(QUAS:mApple-CAAX;he1.1:mCherry)* were injected with *trans*-Tango constructs and Tol2 mRNA at the 1-cell stage. Larvae positive for all secondary markers were imaged by confocal microscopy at 6 dpf. (a) *TgBAC(ptf1a-Gal4-VP16;UAS:GFP)* produced *mApple-CAAX* labeling of hindbrain and midbrain neurons (n= 932/1042). (b) *Tg(th:Gal4-VP16;UAS:GFP)* resulted in extensive labeling throughout the CNS (n= 548/656). (c) In *Tg(-17.6isl2b:Gal4-VP16;myl7:TagRFP;UAS:GFP)* larvae, mApple labeled neurons in the optic tectum (arrowhead; n= 223/297) were adjacent to GFP labeled axon terminals of presynaptic ganglion cells. Diffuse red fluorescent protein labeling (asterisk) is due to a secondary marker (*myl7:RFP*) expressed in the heart. (a’, b’, c’) Merged images with GFP channel. (d, e, f) mApple labeling was not observed after embryos were injected with only the receptor and arrestin constructs (n=25 for each driver). Scale bars, 50 µm.

To address the specificity of *trans*-Tango, we compared the location of mApple labeled neurons with those observed using two other Gal4 driver lines. Cis-regulatory sequences from the *tyrosine hydroxylase* (*th*) gene in *th:Gal4-VP16;UAS:GFP* direct expression in dopaminergic neurons throughout the larval CNS (^26^). Those from the *ISL LIM homeobox 2b* (*islet2b*) gene in *isl2b:Gal4* drive expression in retinal ganglion cells (^27^), as well as in Rohan-Beard sensory neurons, neurons in cranial ganglia, dorsal root ganglia, the anterior and posterior lateral line and populations in the brain (^28–30^). At the 1-cell stage, embryos derived from matings between fish bearing these Gal4 driver lines and a UAS:GFP transgene with fish from the *QUAS:mApple-CAAX* reporter line were injected with the *trans*-Tango ligand, receptor, and arrestin constructs. At 6 days post fertilization (dpf), each driver line produced a distinct pattern of mApple-CAAX labeled neurons and, in all cases, putative postsynaptic cells were located near GFP-labeled neurons or their processes (Figs. 2a,a’, 2 b,b’ and 2c,c’). mApple-CAAX labeled cells were not detected in larvae bearing all *trans*-Tango components except the ligand (Figs. 2d,e,f). These observations indicate that *trans*-Tango functions effectively in the zebrafish nervous system with a high signal-to-noise ratio and patterns of labeling determined by the presynaptic subpopulation that expresses the ligand.

We tested whether transsynaptic labeling could also be achieved with another nearly pan-neural promoter, that of the *Xenopus tubulin beta 2B class IIb* (*Xla.tubb2b*) gene (^31^), which drives expression in the zebrafish CNS from larval to adult stages. Injection of the receptor and arrestin-TEV constructs under *Xla.tubb2b* control together with the ligand plasmid and Tol2 RNA into embryos from matings between the *isl2b:*Gal4 and *QUAS:mApple-*CAAX lines resulted in robust mApple-CAAX labeling as early as 1 dpf, which increased as development proceeded (Fig. 3). Uniquely identified neurons were tracked over time, including commissural primary ascending (CoPA) interneurons, V2a interneurons, and primary motoneurons such as CaP that are all known post-synaptic targets of ligand-expressing Rohon-Beard cells (^32, 33^). Labeling from the *Xla.tubb2b* driven constructs persisted, as mApple positive neurons present in the optic tectum at 6 dpf could still be detected in sections prepared from 2-week-old larvae (Supplemental Fig. 2a-b’).

**Fig. 3:**
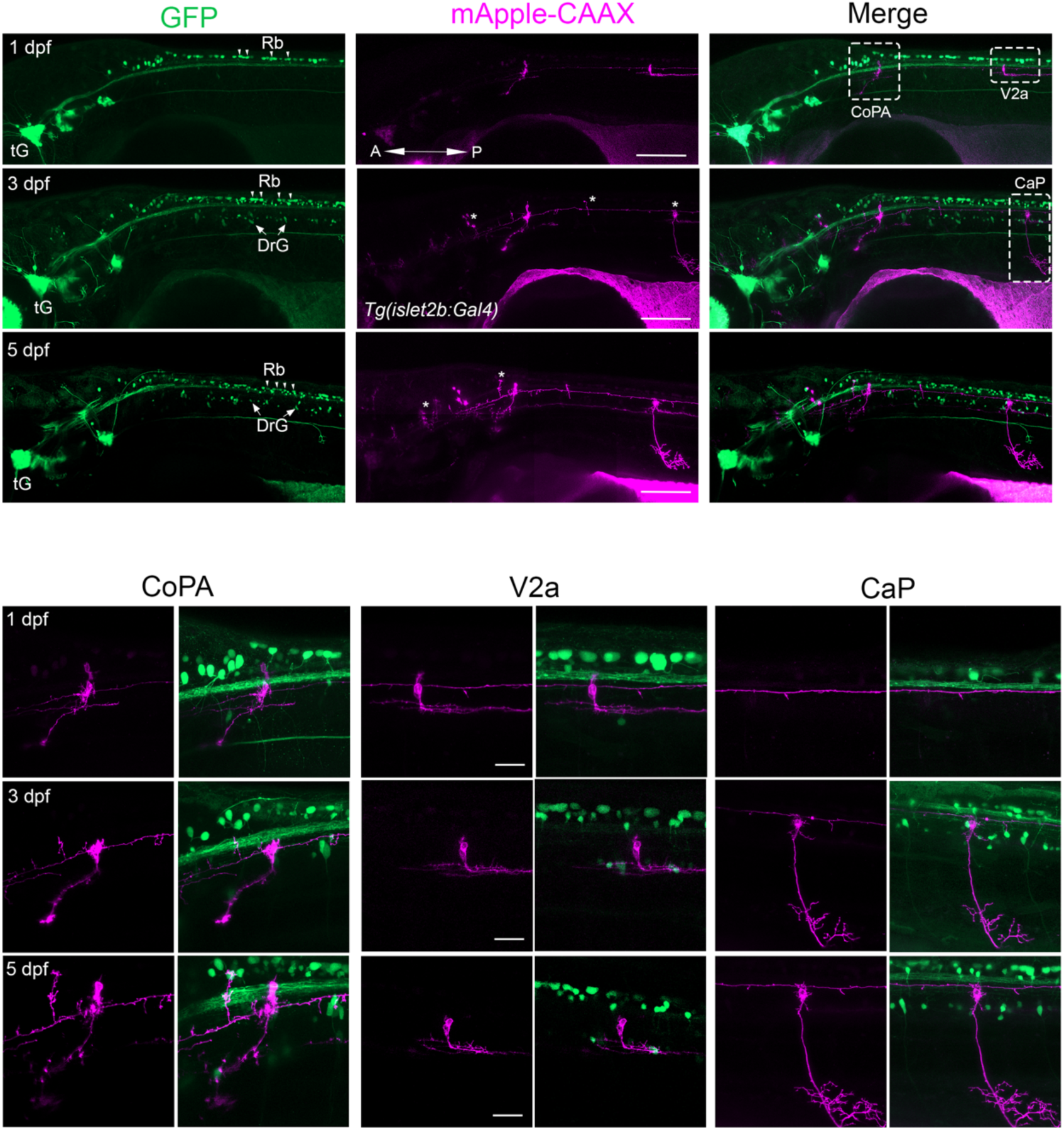
Progression of *trans*-Tango labeling during neural development Lateral views of GFP (pre-synaptic) and mApple-CAAX (post-synaptic) labeled neurons and axons in the ventral spinal cord of the same individual detected at (a) 1 (b) 3 and (c) 5 dpf. The larva contained *Tg(isl2b.2:Gal4; myl7:tagRFP); Tg(UAS:GFP)*; *Tg(QUAS:mApple-CAAX;he1.1:mCherry)* and secondary reporters for all *trans*-Tango components. Additional mApple-CAAX labeled cells were detected over time (arrowheads). Uniquely identified commissural primary ascending (CoPA) interneuron, V2a interneuron and caudal primary motor neuron (CaP) were imaged at each timepoint but, at 1 dpf, the CaP motoneuron was not yet labeled. For each time point, n=10 larvae were examined. Scale bars, 50 µm in upper panels and 100 μm in lower panels.

### Validation of anterograde labeling in the larval retina

The retina is a highly conserved structure that is populated by distinct neuronal cell types distinguished by their morphology, position and connectivity, making it an ideal substrate for testing the fidelity of *trans-* Tango. We identified a total of 102 cells individually labeled by mApple-CAAX in retinal samples from 112 larvae that contained the *pt1fa:Gal4* driver and *QUAS:mApple-CAAX* reporter transgenic lines and all *trans*-Tango plasmids. The *ptf1a* gene is expressed in a retinal progenitor population that gives rise to all amacrine and horizontal cells (^24^). The postsynaptic partners of amacrine cells are retinal ganglion cells and bipolar cells as well as other amacrine cells, whereas horizontal cells form reciprocal synapses with photoreceptors ^34^. Consistent with this, we detected mApple-CAAX labelled cells located in the ganglion cell layer with axons extending to the brain within the optic nerve (Fig. 4a,a’; n=20), in amacrine cells (Fig. 4b,b’; n=35) and bipolar cells (Fig. 4c,c’; n=25) in the inner nuclear layer, and photoreceptor cells in the photoreceptor layer (Fig. 4 d,d’; n=57).

**Fig. 4:**
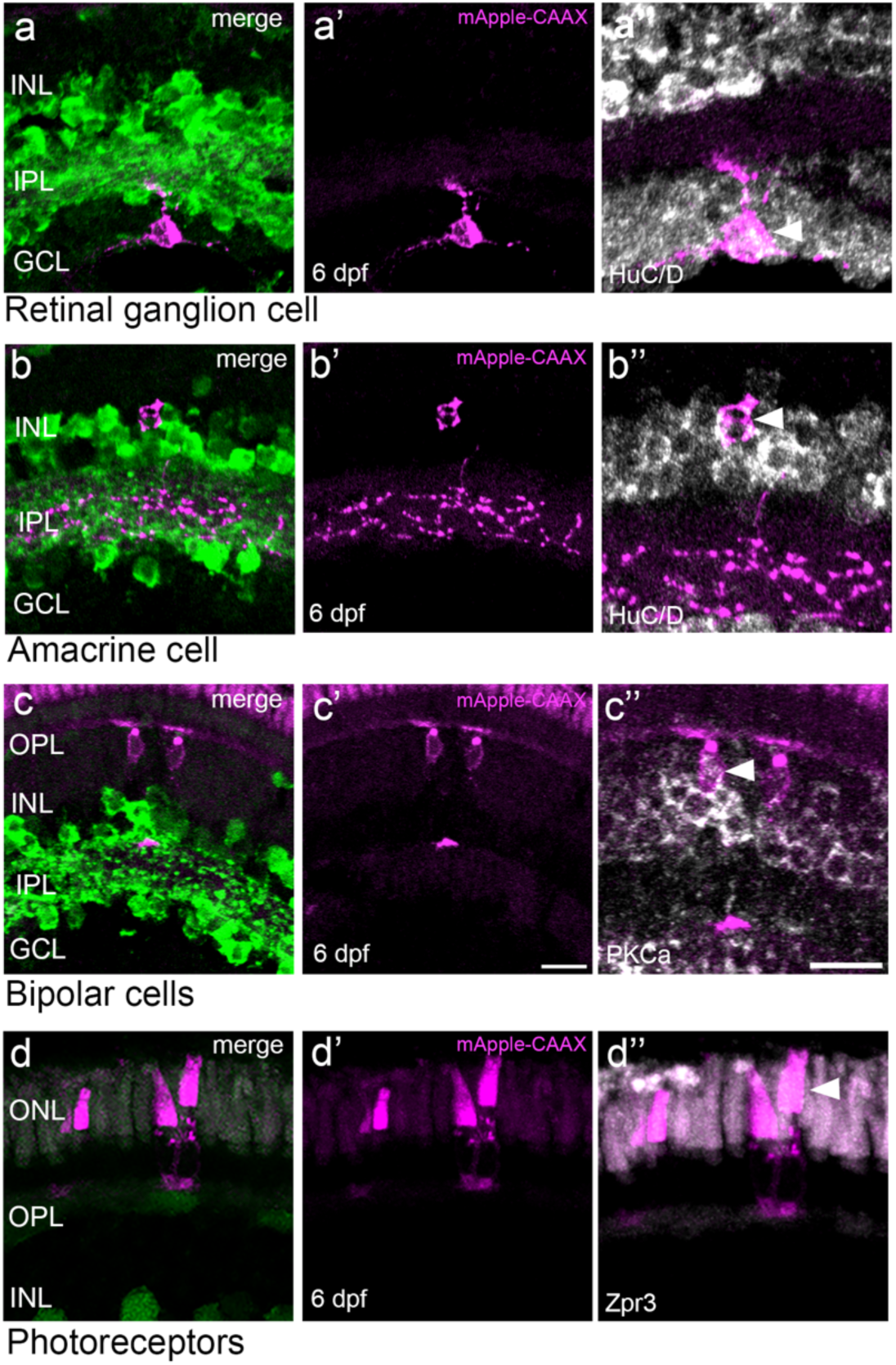
Validation of *trans*-Tango labeling in the retina Co-labeled neurons in cryosections (20 mm) of the retina from *Tg(ptf1a:Gal4-VP16)* and *Tg(QUAS:mApple-CAAX)* larvae that had been injected with *trans*-Tango components at the 1-cell stage. (a,a’) mApple labeled cell in the ganglion cell layer (GCL) with dendrites in the inner plexiform layer (IPL) shows (a’’) HuC/D immunoreactivity (arrowhead). (b,b’) mApple labeled amacrine cell in the inner nuclear layer (INL) with extensive processes in the IPL is also (b’’) HuC/D positive (arrowhead). (c,c’’) mApple labeled bipolar cells in the INL with dendrites in the outer plexiform layer (OPL) variably express protein kinase C alpha (PKCa) (arrowhead). The cone photoreceptor outer segments are visible due to autofluorescence. (d,d’’) Photoreceptors co-labeled with mApple and the zpr-3 monoclonal antibody (ref) are located dorsal to GFP labeled horizontal cells . All retinal sections are shown with photoreceptors oriented at the top. Scale bars, 10 μm.

To corroborate the identities of *trans*-Tango labeled neurons, we performed immunohistochemistry using cell-type specific antibodies and imaged double-labeled cells in cryosections of 6 dpf larval retinas. mApple-CAAX labeled retinal ganglion cells co-labeled with HuC/D antibody were detected in close proximity to GFP-labeled presynaptic amacrine cells (n=8; Fig. 4a’’). Amacrine cells situated in the inner nuclear layer with extensive dendritic processes in the inner plexiform layer also co-labeled with the HuC/D antibody (n=9; Fig. 4b’’), bipolar cells were distinguished by PKCa immunoreactivity (n=6; Fig. 4c’’), and photoreceptors by zpr-3 monoclonal antibody (^35^) (n=; Fig.4d’’).

These results indicate that *trans*-Tango reliably identifies known synaptic partners in the retina.

### *Trans-*Tango in stable transgenic lines

An advantage of plasmid injections is that diffuse labeling enables the identification of individual neurons; however, for some applications, labeling an entire neuronal sub-population is desired. To accomplish this, we established stable lines for each of the *trans*-Tango constructs. Plasmids were injected into 1-cell stage *QUAS:mApple-CAAX* embryos. Those with labeled hatching gland cells at 24 hpf were raised to adulthood and their progeny screened to recover transgenic founders with germ-line clones. Founders were mated with *QUAS:mApple-CAAX* fish and their heterozygous progeny used to establish stable lines. Wholemount in situ hybridization (WISH) was performed at 6 dpf to verify transgene expression with RNA probes designed to capture unique sequences only found in the *trans* Tango component (refer to Supplementary Fig. 3). For the ligand line *10xUAS-E1B:sGCG-ICAM(1235) nrxn1b*, the probe detected transcripts in the same pattern as endogenous *ptf1a* expression (^25^), but not in the absence of the *pt1fa:Gal4-VP16* transgene (Supplementary Fig 3a). The arrestin-TEV and receptor transgenes were broadly expressed throughout the CNS (Supplementary Figs. 3b,c) similar to endogenous *elavl3* transcripts (^18^). Sibling embryos that lacked YFP labeling in the hatching gland at 1 dpf did not show any WISH signal as expected (Supplementary Fig. 3b,c).

To confirm that the stable ligand line functioned effectively at transsynaptic signaling, we mated the *pt1fa-*Gal4 driver line with F1 fish bearing *QUAS:mApple-CAAX* and *10xUAS-E1B:sGCG ICAM(1235)-nrxn1b* and injected the receptor and arrestin-TEV constructs into their 1-cell stage embryonic progeny. At 6 dpf, over 80% of larvae (n=257/312) displayed mApple-CAAX labeled postsynaptic neurons (Supplementary Fig. 3d-d’’). We observed similar results with *elavl3:hArr-TEV* larvae that, as embryos, had been injected with the ligand and receptor plasmids (n=132/164; Supplementary Fig. 3e-e’’). Only larvae from the receptor line, *elavl:hGCGR-TEVcs-QF*, failed to produce transsynaptic labeling following injection of the ligand and arresti-TEV constructs. This is unlikely due to transgene position effects as experiments were performed on three independently established transgenic lines, whose larvae showed robust hGCGR-QF expression, but no mApple-CAAX labeled neurons (n= 0/276; data not shown). Moreover, mApple-CAAX neurons were not observed using a stable line for a receptor construct that was codon optimized for zebrafish (hGCGR*) and fused to QF (n= 0/171; Supplementary Fig. 3f,f’,f’’). All transgenic lines did produce functional receptor because robust mApple-CAAX labeling was observed at 6 dpf after a subthreshold concentration of the receptor plasmid (Supplementary Fig. 3g,g’) was injected into *elavl:hGCGR*-TEVcs-QF* embryos (n=21/25; Supplementary Fig. 3h,h’). along with the ligand and arrestin-TEV constructs.

We further examined transsynaptic labeling using the ligand and arrestin-TEV transgenic lines, either individually or together, with a different Gal4 driver, *isl2b:Gal4-VP16*, in which individual neurons could be more easily resolved (Supplementary Fig. 4). Notably, on average, the number of mApple CAAX labeled neurons was significantly greater (>5-fold) after the receptor was introduced into embryos doubly transgenic for both the ligand and arrestin-TEV (Supplementary Fig. 4d,d’ and f) compared to the number observed following injections of all *trans*-Tango plasmids (Supplementary Fig. 4a,a’) or injections into embryos with either the ligand (Supplementary Fig. 4b,b’) or arrestin-TEV transgene alone (Supplementary Fig. 4c,c’). Image registration of ten *isl2b:Gal4-VP16*; *QUAS:mApple-CAAX* 6 dpf larvae bearing the ligand and arrestin-TEV transgenes and injected receptor plasmid revealed overlap in the location of *trans-Tango* labeled postsynaptic neurons (Supplementary Fig. 4g).

### *trans*-Tango validates retinotectal circuitry

Calcium imaging and sparse labeling experiments suggest that tectal neurons with distinct morphologies are the targets of retinal ganglion cells (^36, 37^). To test this connectivity directly, we used the *isl2b:Gal4 VP16* driver line in fish containing *UAS:GFP* and *QUAS:mApple-CAAX* to express *trans*-Tango ligand selectively in retinal ganglion cells. We injected the ligand, receptor and arrestin-TEV constructs with Tol2 mRNA in 1-cell stage embryos, sorted out those having all secondary markers at 1 and 2 dpf, and then screened for *mApple-CAAX*-labeled tectal neurons at 6 dpf.

Seven neuronal subtypes previously proposed as postsynaptic to the optic tectum were uniquely identifiable based on their position and shape (Fig. 5a-i’). These included non-stratified periventricular neurons (nsPVIN, n=46; Fig. 5b,b’), superficial interneurons (SIN, n=11; Fig. 5c,c’), mono-stratified periventricular neurons (msPVIN, n=67; Fig 5d,d’), bi-stratified periventricular interneurons (bsPVIN, n=43; Fig. 5e-e’), periventricular projection neurons (PVPN, n=21; Fig. 5f,f’ and h,h’) and tri-stratified periventricular interneurons (tsPVIN, n=72; Fig. 5g,g’). Using Fiji software, we produced renderings of individually identified cells (Fig 5i) whose morphology closely corresponded to the previously described tectal neurons (^37, 38^).

**Fig. 5.**
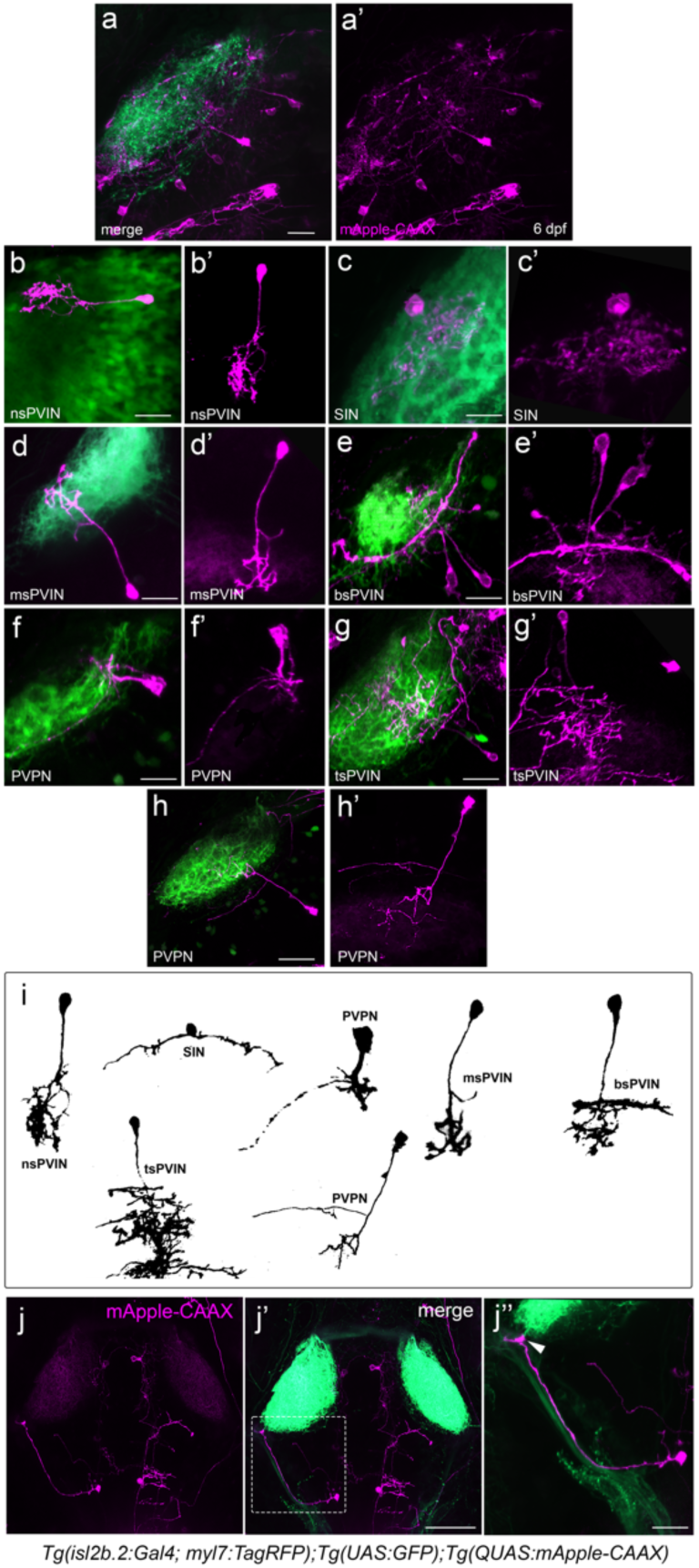
*trans-*Tango confirms predicted postsynaptic partners of retinal ganglion cells (a,a’) *Tg(isl2b.2:Gal4; myl7:tagRFP)* drives UAS:GFP in retinal ganglion cells that project to the optic tectum, where labeling from *Tg(QUAS:mApple-CAAX)* reveals their postsynaptic partners at 6 dpf. (b) Distinct morphology of non-stratified periventricular interneurons (nsPVN, n=46), (c) superficial interneurons (SIN, n=11), (d) mono-stratified periventricular interneurons (msPVIN, n=67), (e-e’) bistratified periventricular interneurons (bsPVIN, n=43) (f) periventricular projection neurons (PVPN, n=21), (g) tri-stratified periventricular interneurons (tsPVIN, n=72) and (h) a distinct subtype of periventricular projection neurons (PVPN, n=26) labeled by *trans*-Tango. In (a’-h’), individual neurons are reoriented with cell bodies anterior. Scale bars = 100 µm. (i) Drawings corresponding to each neuronal cell type. (j-j’) Occasionally, hindbrain neurons with long processes extending to the tectum are also detected (n=6/297). Scale bar 50 μm in j’ and 100 μm in j’’.

With *trans-*Tango we were also able to identify a postsynaptic neuron that exhibited a previously undescribed morphology characterized by a long process in contact with the axon terminals of retinal ganglion cells and a cell body located in the hindbrain (n=6; Fig. 5j-j’’). These data indicate that *trans* Tango consistently labels predicted synaptic partners and is also a powerful tool for revealing unsuspected postsynaptic targets.

### Activation of synaptically-coupled neurons

To verify that the *trans*-Tango labeling correlates with functional synaptic connections, we optogenetically activated retinal ganglion cells in 6 dpf *isl2b:Gal4-VP16* larvae and assessed the response in putative postsynaptic neurons in the optic tectum. Activity in post-synaptic cells was monitored with a nuclear-tagged genetically encoded calcium indicator (H2B-GcaMP6s) (^39^) that we placed under QUAS control. Receptor, arrestin-TEV and QUAS:H2B-GcaMP6s plasmids were injected into the 1-cell stage progeny of *isl2b:Gal4-VP16; QUAS:mApple-CAAX* fish mated to the *10xUAS-E1B:sGCG-ICAM(1235) nrxn1b* ligand line and later screened for secondary markers for all *trans*-Tango components. We first confirmed that GcaMP6s positive nuclei were detected in mApple-CAAX neurons in the 6 dpf larval tectum (dashed boxes, Fig 6a-a’’), indicating that both reporter genes were co-expressed through QF transcriptional regulation. Next, we activated retinal ganglion cells using the same Gal4 line to drive expression of the redshifted channel rhodopsin ReaChR under UAS control (*UAS:ReaChR-Tag-RFPT*) (40) and recorded the change in fluorescence (ΔF/F) of GcaMP6s positive tectal neurons following retinal exposure to a 561 nm laser. Individual neurons were imaged in the region of the tectum where axon terminals from retinal ganglion cells form a layered neuropil and connect to dendrites and axons of interneurons and projection neurons (^37, 41^). We observed a significant increase in calcium transients in larvae containing *UAS:ReaChR-Tag-RFPT* compared to sibling larvae that lacked this transgene (p= 0.0048; Fig. 6b-d). Optogenetic activation of retinal ganglion cells, thus, leads to a response in tectal neurons that is visualized through GcaMP6s, the product of *trans*-Tango transsynaptic signaling.

**Figure 6.**
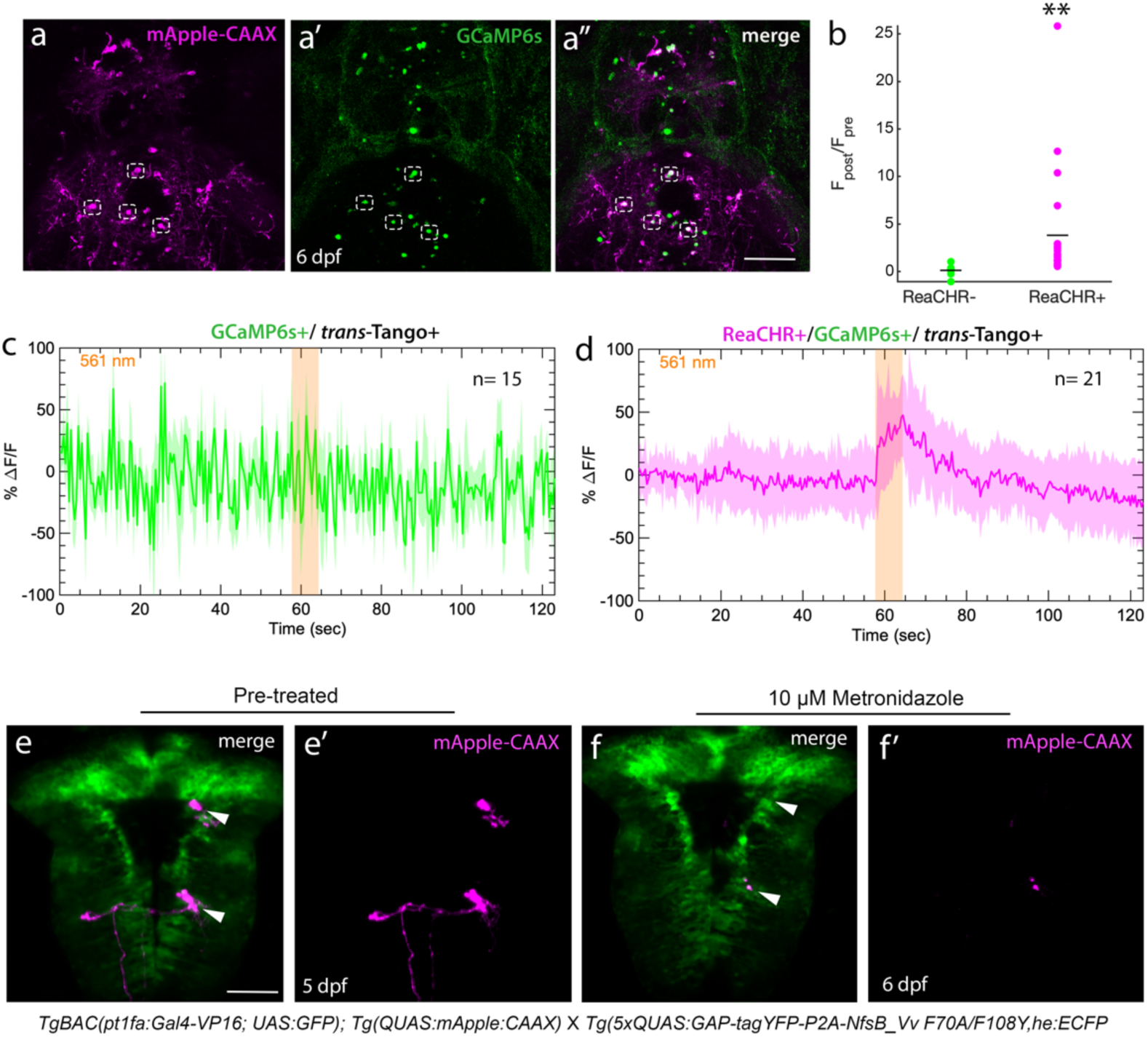
Functional validation of synaptic connectivity *trans*-Tango signaling drives expression of any gene under QUAS control. (a,a’’) In *Tg(isl2b.2:Gal4; myl7:tagRFP)*, *Tg(UAS:GFP)* and *Tg(QUAS:mApple-CAAX)* larvae that were injected with the QUAS:H2B-GCaMP6s plasmid, GCaMP6s labeled nuclei are only detected within mApple-CAAX labeled tectal neurons of *trans*-Tango positive larvae. (b-d) In larvae bearing all *trans*-Tango components, optogenetic activation of retinal ganglion cells expressing In larvae bearing all *trans*Tango components, optogenetic activation of retinal ganglion cells expressing ReaChR with 561 nm light activates GCaMP6s in tectal neurons. (b) Graphical representation showing 5-fold increase in calcium transients in *Tg(UAS:ReaChR-Tag-RFPT)* larvae compared to siblings lacking this transgene (p = 0.0043). (c-d) Mean and standard deviation of ΔF/F for GCaMP6s positive tectal neurons in (c) ReaChR negative controls (n=15) compared to (d) ReaChR positive larvae (n=21) in response to 561 nm of light. (e,e’) *trans*-Tango mediated ablation of postsynaptic neurons. In 5 dpf *TgBAC(ptf1a:Gal4-VP16)*^jh16^; UAS:GFP) larvae expressing the bacterial *NfsB* reducatase gene, mApple-CAAX neurons (arrowheads) were no longer detected (f,f’) following 1 day of incubation in 10 μM Metronidazole (n= 25). Scale bars, 50 μm.

Another demonstration of *trans*-Tango mediated gene regulation is the targeted ablation of postsynaptic neurons. We mated fish from the ptf1a:Gal4 driver line containing the *QUAS:mApple-CAAX* reporter to a transgenic line that has an optimized version of the bacterial nitroreductase gene *nfsB* under QUAS control *5xQUAS:GAP-tagYFP-P2A-NfsB_Vv F70A/F108Y* (^42^). At 5 dpf, larvae were imaged for *mApple-CAAX* labeling in the hindbrain (Fig. 6e,e’) and then incubated for 24 hours in 10 mM of metronidazole, which when reduced is toxic to cells. Reimaging of the same hindbrain region revealed that *mApple-CAAX* neurons were no longer detected (Fig. 6f-f’). The results indicate that *trans*-Tango signaling can be used to identify, monitor, and disrupt neural circuits in the zebrafish brain.

## Discussion

We established an anterograde transsynaptic labeling system, *trans*-Tango, for the zebrafish CNS. This is the first genetically encoded system for transsynaptic labeling in a vertebrate nervous system. We used *trans*-Tango to regulate the expression of transgenic reporters in the postsynaptic partners of genetically identified presynaptic neurons. These reporters can be used to trace connectivity anatomically and to identify postsynaptic partners of the starter neurons as well as monitor their activity or ablate them.

Functional investigations into neural pathways in zebrafish have been facilitated by optogenetic modulators and genetically encoded calcium indicators that report neuronal responses throughout the larval brain (^43–45^). However, genetic tools to validate synaptic connections directly have been lacking. Retrograde tracing of afferents using recombinant rabies, vesicular stomatitis, or herpes simplex 1 virus has been applied to the zebrafish CNS with some success (^46, 47^). However, drawbacks to viral approaches include the potential for neuronal toxicity, the necessity to inject recombinant virus locally into brain regions, and the maintenance of larval fish at higher than usual temperatures for effective expression.

Unlike the available tools for circuit interrogation in zebrafish, the *trans*-Tango genetic strategy is solely based on the introduction of an optimized set of plasmids into 1-cell stage embryos. Hundreds of embryos can be readily injected, providing a large sample size for testing constructs. Moreover, concentration levels can be adjusted to recover broad or sparse labeling for assessing connectivity between neuronal populations or single synaptic partners, respectively. Expression of the *trans*-Tango ligand under UAS transcriptional control capitalizes on the large repertoire of neural-specific, transgenic Gal4 driver lines available in zebrafish (^38, 48–52^), while plasmid injections avoid the transcriptional silencing often associated with UAS-regulated transgenes in the zebrafish genome across generations (^53–55^).

A major advantage of the *trans*-Tango approach is the ability to perform live imaging to monitor neural connections in the same individual over time, as was demonstrated for postsynaptic targets of *isl2b* neurons. With persistently expressing pan-neural promoters, *trans*-Tango can also be applied to examine circuits in the juvenile and adult CNS. Ideally, performing analyses on adult brains will benefit from providing *trans*-Tango reagents as stable transgenes, as in *Drosophila* (^7^). Although we were successful at accomplishing this with transgenes for the ligand and arrestin-TEV components, we have not yet achieved *trans*-Tango labeling using several independently isolated receptor transgenic lines. Similar negative results were obtained when hGCGR or codon-optimized hGCGR* constructs were driven by either the *elavl3* or *Xla.tubb2b* promoter. One explanation is that an insufficient number of receptors are concentrated at postsynaptic sites, owing to their distribution throughout the extensive membranes of neuronal processes. In support of this, individuals bearing four copies of receptor transgenes that had been injected with ligand and arrestin-TEV plasmids did not exhibit labeling of the postsynaptic reporter (data not shown). However, robust postsynaptic labeling was observed when a concentration of receptor plasmid, that itself is too low to mediate *trans*-Tango signaling, was injected into animals that had only one copy of the receptor transgene. These findings indicate that transgenic lines indeed produce functional hGCGR. We further determined that injection of the hGCGR-QF construct into embryos bearing both the ligand and arrestin transgenes, yielded significantly more labeled neurons compared to injections into each transgenic line individually. Experimental approaches requiring widespread transsynaptic signaling can therefore rely on receptors derived from plasmid injected into doubly transgenic animals homozygous for the ligand and arrestin-TEV transgenes.

Distinct patterns of labeling were obtained using three different Gal4 driver lines. In all cases, postsynaptic neurons demarcated by the QUAS-controlled membrane-tagged reporter were in close proximity to neurons or their processes expressing the UAS-regulated reporter. Image registration also revealed an overlap in the positions of labeled cells between individual larvae from a given Gal4 driver line. To assess the fidelity of connections between neurons, we turned to the retina where there is considerable information about the synaptic relationships between cell types (^34^). As expected, based on their position and immunolabeling, horizontal cells expressing ligand yielded *trans*-Tango labeling of bipolar cells, whereas ligand producing amacrine resulted in labeling of other amacrine cells or retinal ganglion cells. Ganglion cells are the only retinal neurons that project to the brain via the optic nerve, where they terminate in discrete arborization fields in the optic tectum (^56, 57^). Prior studies in several fish species revealed the stereotypic morphology of defined classes of tectal neurons from Golgi labeling, application of dyes, or calcium imaging of neural activity (^36, 37, 58^). Remarkably, by directing *trans*-Tango in retinal ganglion cells, we identified the same classes of neuronal subtypes that were previously reported in the tectum, supporting the validity of this genetic approach to confirm postsynaptic targets with high specificity. We also observed connectivity with a previously undescribed cell type that extends a long process from its cell body situated in the hindbrain. These findings further demonstrate the potential of *trans*-Tango to reveal new synaptic partners.

An important feature of a genetic, anterograde transsynaptic signaling pathway is the genetic access that it provides to postsynaptic neurons. With *trans-*Tango, expression is dependent on translocation of the QF transcription factor from the cell membrane to the nucleus where it promotes transcription of any gene under QUAS control, either from plasmids or genomic transgenes. Increasing numbers of QUAS driven reporter and effector transgenic lines are available in zebrafish (^15, 59–62^) and all can be used as outputs of *trans-*Tango signaling. In addition to the membrane-tagged fluorescent reporter mApple-CAAX, we present functional evidence for activation of QUAS regulated genes encoding a nuclear GcaMP and a bacterial nitroreductase. Together, such *trans*-Tango dependent tools offer new ways to characterize synaptically coupled neurons and evaluate how their activity within neural circuits underlies behavioral responses. Implementation in zebrafish of *retro*-Tango, a recently described Tango-based strategy for retrograde transsynaptic signaling (^63^), will further increase the versatility and power of genetic approaches to probe neural pathways both acutely and over time in a vertebrate brain.

## Methods

### Animals

Zebrafish were reared under a 14:10 light:dark cycle at 27°C in dechlorinated and filtered system water. Transgenic lines for *trans-*Tango components were generated in the AB wild-type strain (^64^) including two alleles of *Tg(10xUAS-E1B:sGCG-ICAM(1235)-nrxn1b; cryaa:mCherry)*^cd^^28, cd^^29^ referred to as (*10xUAS-E1B:sGCG-ICAM(1235)-nrxn1b)*, four alleles of *Tg(elavl3:hGCGR-TEVcs-QF; he1.1:CFP)*^c796, c797, cd30, cd34^ (*elavl3:hGCGR-TEVcs-QF*) and *Tg(elavl3:hArr-TEV)*^cd^^30^ (*elavl3:hArr-TEV)*. The Gal4 driver lines *TgBAC(pt1fa:Gal4-VP16; UAS:GFP)^jh^*^16^ (*pt1fa:Gal4-VP16*) (^65^), *Tg(th:Gal4-VP16; UAS:GFP)^m^*^1233^ (*th:Gal4-VP16)* (^66^) and *Tg(-17.6isl2b:Gal4-VP16;myl7:RFP;UAS:GFP)*^zc^^60^ (*isl2b:Gal4-VP16)* (^67^). And QUAS regulated lines *Tg(QUAS:mApple-CAAX; he1.1:mCherry)^c^*^6^^36^ (*QUAS:mApple-CAAX*)^68^ and *Tg(5xQUAS:GAP-tagYFP-P2A-NfsB_Vv F70A/F108Y,he:ECFP)^jh5^*^62^ (^42^) were also used in this study. To inhibit melanin pigmentation, embryos and larvae were maintained in system water containing 0.003% phenylthiourea (PTU; P7629, Sigma-Aldrich). All protocols and procedures were approved by the Institutional Animal Care and Use Committee (IACUC) of Dartmouth College.

### Plasmids

Plasmids for the *trans*-Tango components were generated using the Hi-Fi DNA Assembly (NEB #E5520S), BP Gateway Cloning (Thermo Fisher, Invitrogen #11789020) and LR Gateway Cloning (Thermo Fisher, Invitrogen #11791020) kits. Relevant PCR primers indicated below are listed in Table 1. All ligand constructs were generated in a similar manner. The plasmid pExpTol2-UAS-E1B-ReaChR-TS-tagRFP-cryaa-mCherry was generated as an intermediate construct. The cryaa-mCherry sequence was amplified from pTol2pA2-ubb:secHsa.ANNEXINV-mVenus, cryaa:RFP (Addgene #92388) using primers 3283 and 3284. The SV40 late polyA sequence was amplified from pKHR8 (Addgene #74625) using primers 3285 and 3286. The pExpTol2-UAS-E1B-ReaChR-TS-tagRFP plasmid was digested with AscI and MluI. Both PCR products and the digested vector were assembled using Hi-Fi DNA Assembly to obtain pExpTol2-UAS-E1B-ReaChR-TS-tagRFP-cryaa-mCherry. For pExpTol2-UAS-E1B-sGCG-Nrxn1a; cryaa-mCherry, the sGCG sequence was amplified from the *Drosophila trans*-Tango plasmid (^7^) using primers 3274 and 3279. To create the pExpTol2-UAS-E1B-sGCG-ICAM(1235)-Nrxn1a; cryaa mCherry vector, sGCG-ICAM(1235) sequences were amplified from the *Drosophila* plasmid (^7^) in two separate reactions, using primer pairs 3274 and 2302 and 3275 and 2303. For pExpTol2-UAS-E1B-sGCG-ICAM-Nrxn1a; cryaa-mCherry the sGCG-ICAM sequence was amplified using the primers 3274 and 3275. Nrxn1a sequence encoding the full-length protein without the signal peptide was amplified using primers 3280 and 3264. Sequences encoding the transmembrane and the intracellular domains of Nrxn1a swere amplified using primers 3276 and 3264 from pooled cDNA of 6 dpf zebrafish larvae. PCR products were subsequently cloned pExpTol2-UAS-E1B-ReaChR-TS-tagRFP-cryaa-mCherry (Addgene #43963) digested with SpeI and PacI using Hi-Fi DNA Assembly.

**Table 1:**
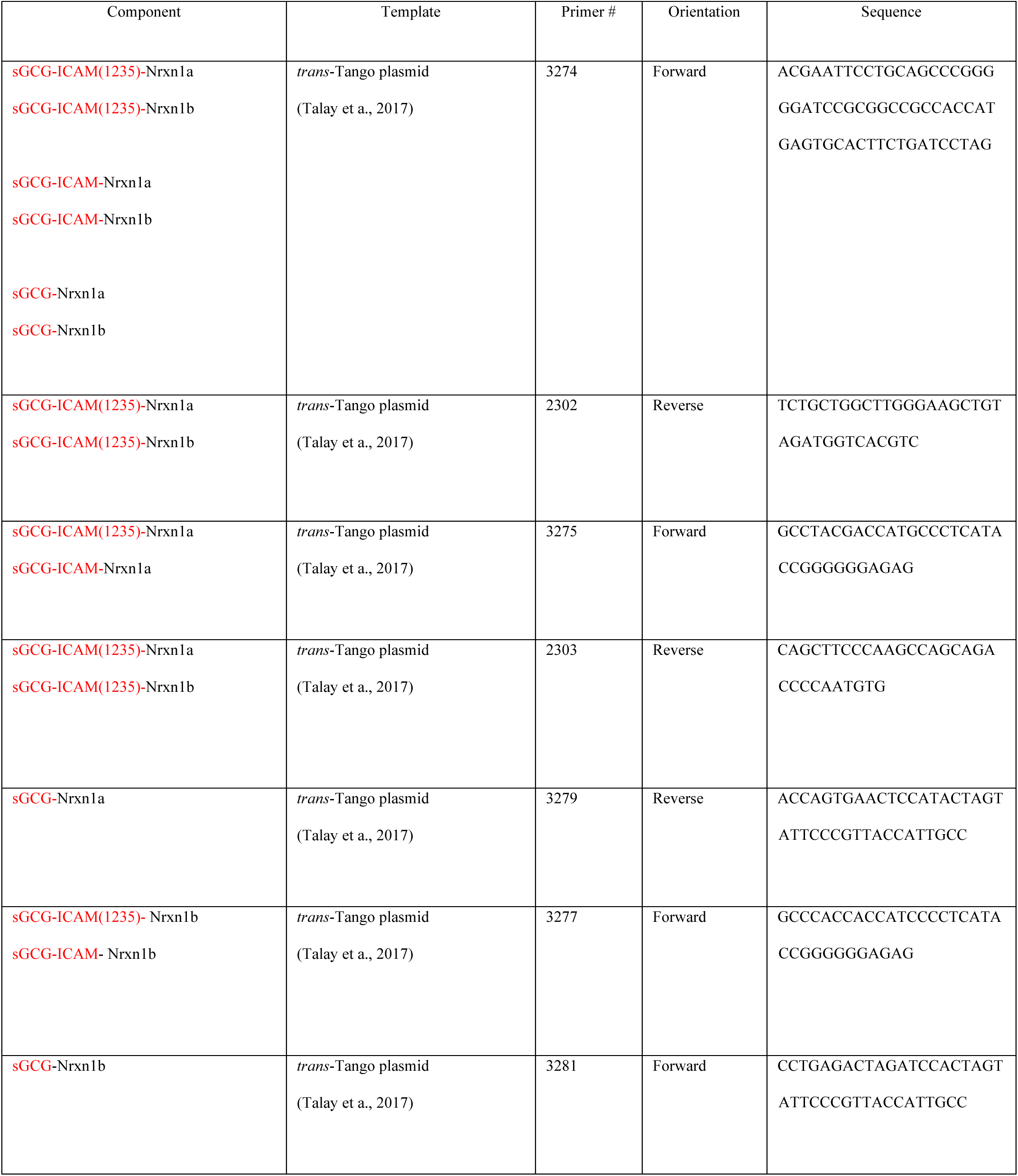
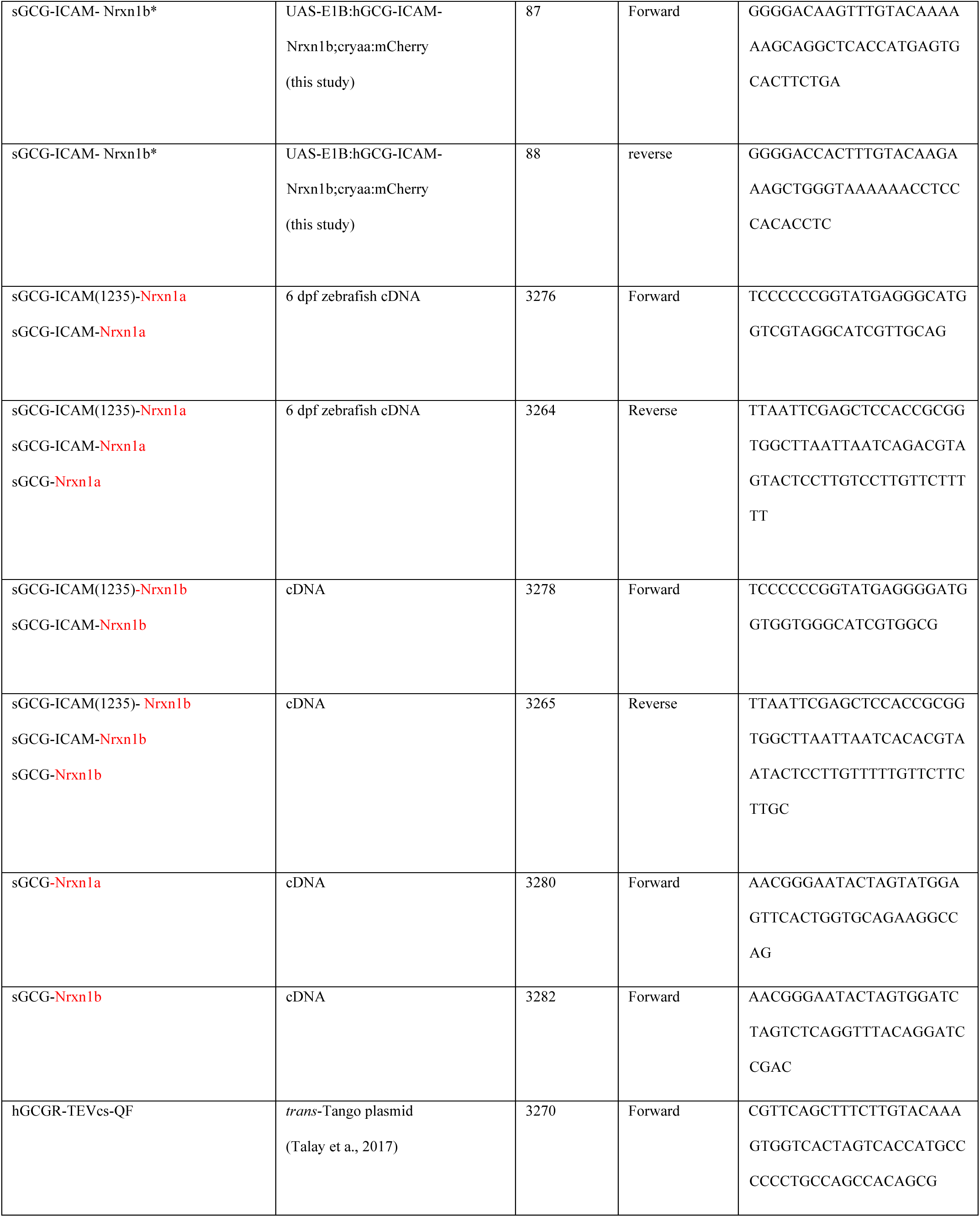
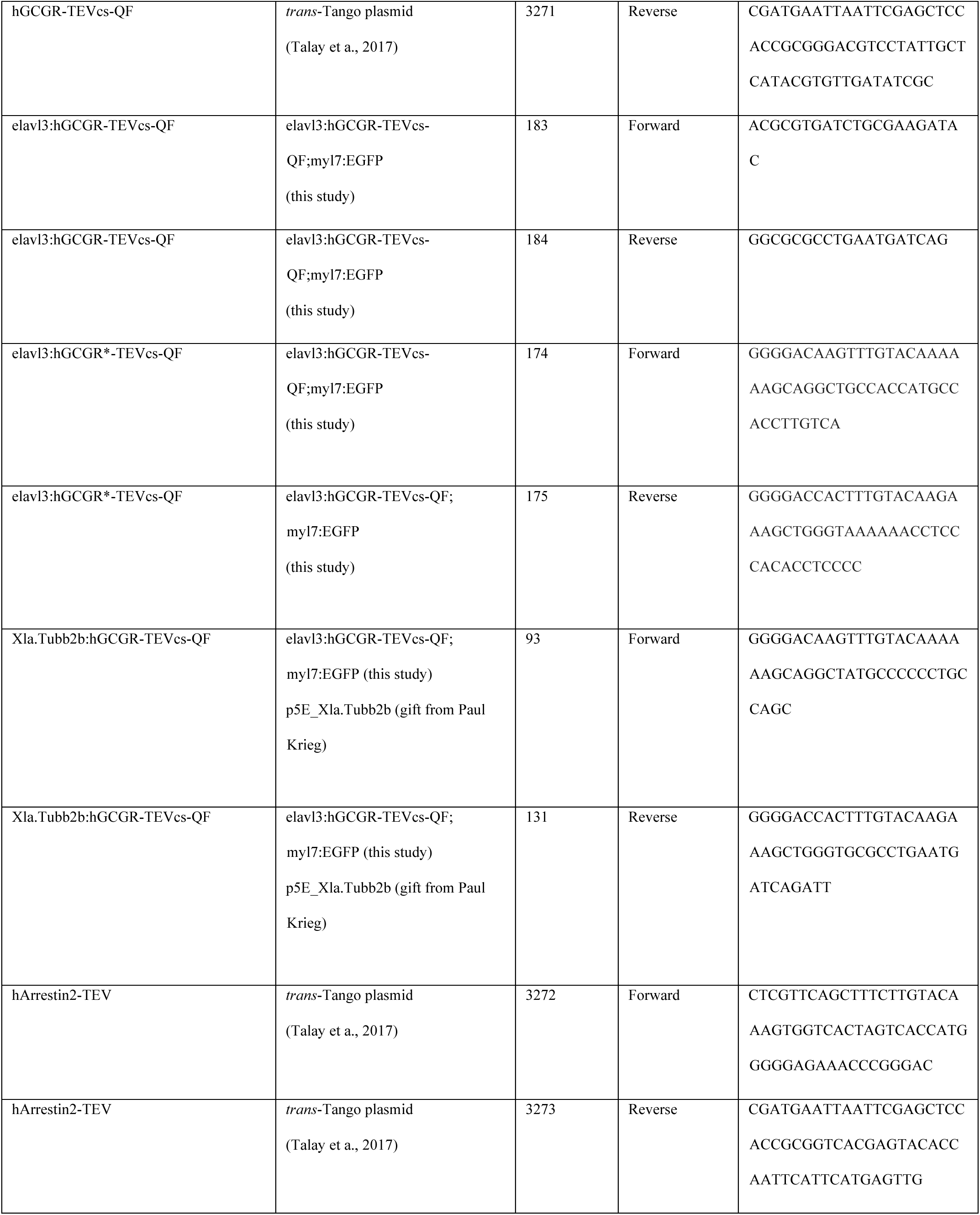
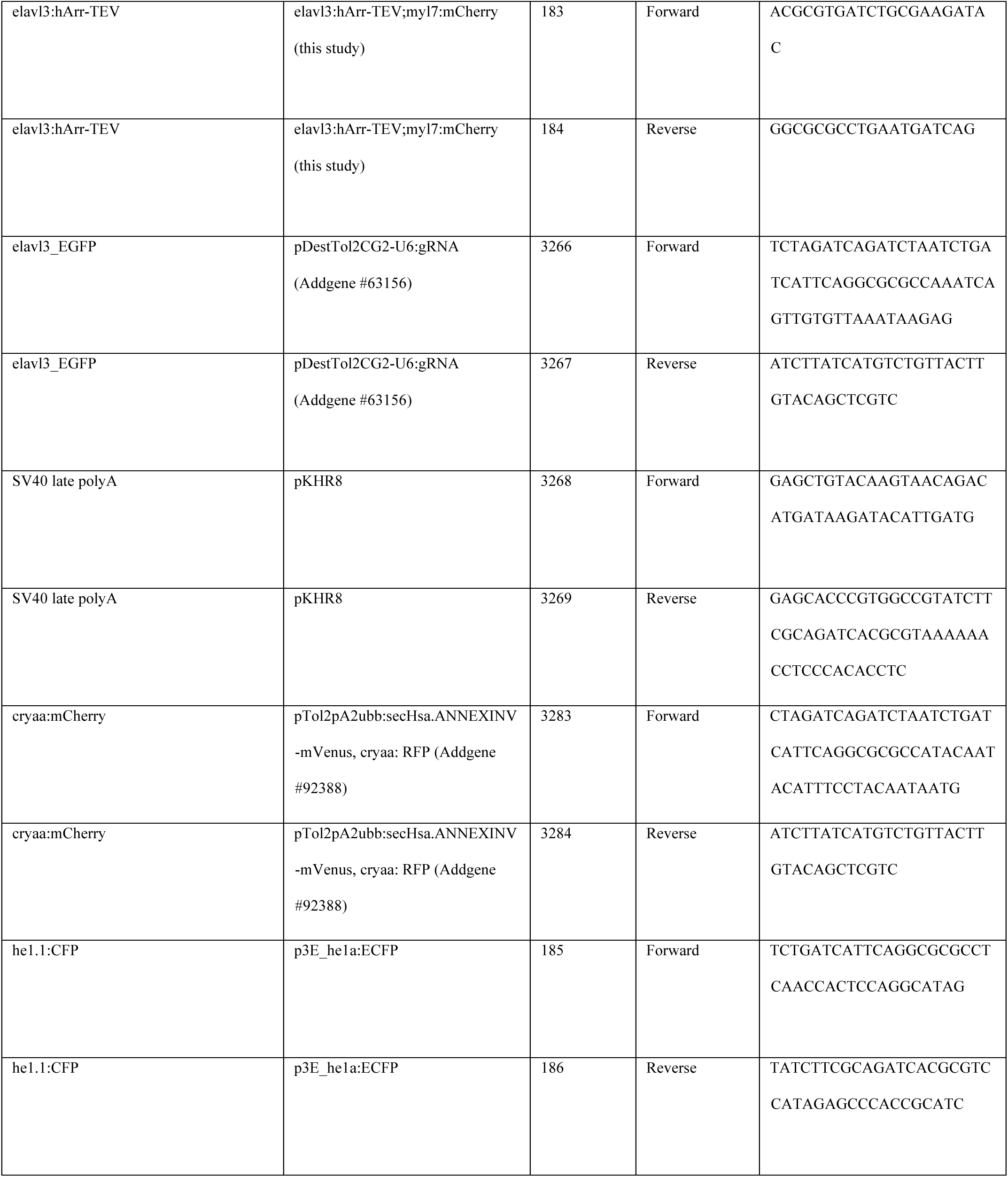
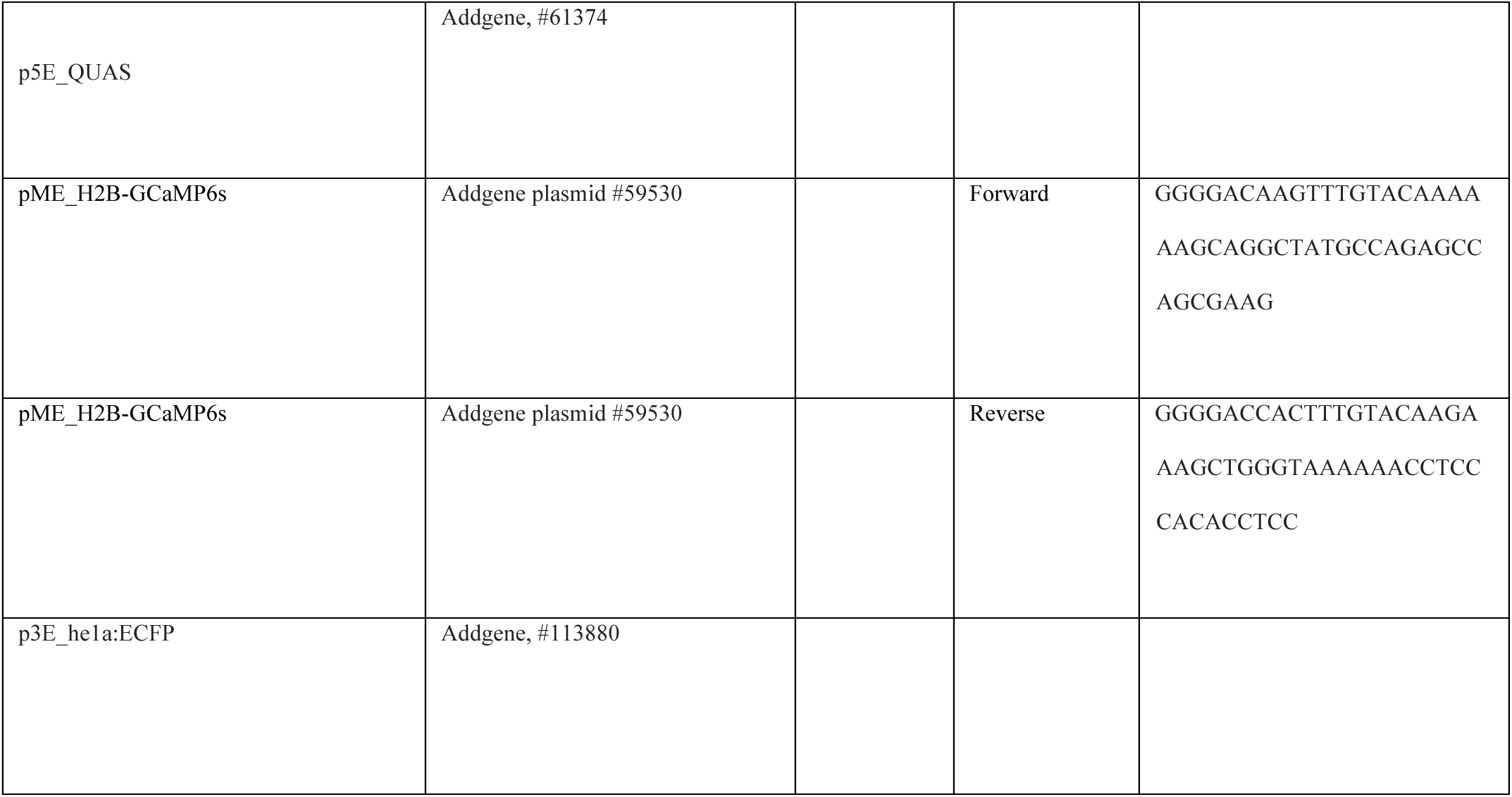
Primers for synthesis of plasmids. PCR amplified components of plasmids are indicated by the text in red.

Constructs for the Nrxn1b plasmids were generated as described above. For pExpTol2-UAS:E1B-sGCG, the sGCG sequence was amplified from the trans-Tango plasmid (^7^) using primers 3274 and 3281. To create pExpTol2-UAS-E1B-sGCG-ICAM(1235)-Nrxn1b; cryaa-mCherry, the sGCG-ICAM(1235) sequence was amplified from the trans-Tango plasmid (^7^) in two separate reactions using the primer pairs 3274 and 2302 and 3277 and 2303. For pExpTol2-UAS-E1B-sGCG-ICAM-Nrxn1b; cryaa-mCherry, the sGCG-ICAM sequence was amplified from the *trans*-Tango plasmid (^7^) using primers 3274 and 3277.

Nrxn1b sequence encoding full-length protein without the signal peptide was amplified from zebrafish cDNA using primers 3282 and 3265 and the transmembrane and the intracellular domains were amplified using primers 3278 and 3264. PCR products were subsequently cloned into pExpTol2-UAS-E1B-ReaChR-TS-tagRFP-cryaa-mCherry (Addgene #43963) digested with SpeI and PacI using Hi-Fi DNA Assembly.

To produce hGCGR vectors, myl7_EGFP sequence was amplified from pDestTol2CG2-U6:gRNA (Addgene #63156) using primers 3266 and 3267. The SV40 late polyA sequence was amplified from pKHR8 (Addgene #74625) using primers 3268 and 3269. The Tol2 Gateway Destination Vector (^43^) was digested with AscI and MluI. Both PCR products and the digested vector were assembled using Hi-Fi DNA Assembly. The resulting plasmid was subsequently digested with SpeI and SacII. The hGCGR-TEVcs-QF sequence was amplified from the *Drosophila trans*-Tango plasmid (^7^) using primers 3270 and 3271. The PCR product and the digested vector were assembled using Hi-Fi DNA Assembly. The final plasmid was then subjected to LR Gateway cloning with pEntry-HuC (^43^) to obtain elavl3:hGCGR-TEVcs-QF; myl7:EGFP. In later experiments, the secondary reporter labeling the heart (myl7:EGFP) was replaced with he1.1:CFP that labels hatching gland cells. The he1.1:CFP sequence was amplified from p3Ehe1a:ECFP (Addgene #113880) using primers 185 and 186.

The receptor construct was also placed under the control of the Xla.Tubb2b promoter (Xla.Tubb2b:hGCGR-TEVcs-QF; he1.1:CFP). The hGCGR-TEVcs-QF sequence was amplified from elavl3:hGCGR-TEVcs-QF;myl7:EGFP using primers 93 and 131 that contained flanking attB1 and attB2 sequences, respectively. A Gateway recombination reaction was performed using BP Clonase II with this PCR product and donor vector pDONR221 (Invitrogen, #12536017) to obtain pME_hGCGR-TEVcs-QF. Using Gateway LR Clonase II, p5E nbt (^59^), pME_hGCGR-TEVcs-QF, p3E_he1a:ECFP (Addgene #113880) were introduced into the Tol2 Gateway Destination Vector (^69^) (#394) to obtain Xla.Tubb2b:hGCGR-TEVcs-QF;he1.1:CFP.

The hGCGR sequence from the elavl3:hGCGR-TEVcs-QF;myl7:EGFP (this study) was codon optimized for zebrafish (hGCGR*) following https://www.nichd.nih.gov/research/atNICHD/Investigators/burgess/software (^70^) and synthesized by Gene Universal (www.geneuniversal.com) into pUC57-BsaI. From this plasmid, the hGCGR* sequence was amplified and attB1 and attB2 arms attached using primers 174 and 175. The PCR product was placed in pME (pDONR221 #398, Addgene) using a Gateway recombination reaction with BP clonase II. Subsequently, Gateway reactions were used to introduce either p5E_elavl3 or p5E_nbt with pME_hGCGR*-TEVcs-QF and p3E_he1a:ECFP (Addgene #113880) into the Tol2 Gateway Destination Vector (^69^) (#394) to produce elavl3:hGCGR*-TEVcs-QF;he1.1:CFP or Xla.Tubb2b:hGCGR*-TEVcs-QF;he1.1:CFP.

For the arrestin construct elavl3:hArr-TEV;myl7:mCherry, the myl7:mCherry sequence was amplified from pKHR8 (Addgene #74625) using primers 3266 and 3267. The SV40 late polyA sequence was amplified from pKHR8 using primers 3268 and 3269. A Tol2 Gateway Destination Vector (Ahrens et al., 2012) was digested with AscI and MluI. Both PCR products and the digested vector were assembled using Hi-Fi DNA Assembly. The resulting plasmid was subsequently digested with SpeI and SacII. The hArrestin2-TEV sequence was amplified from the trans-Tango plasmid (^7^) using primers 3272 and 3273. The PCR product and the digested vector were assembled using Hi-Fi DNA Assembly. The final plasmid was then subjected to LR Gateway cloning with pEntry-HuC (^43^) to obtain elavl3:hArr-TEV; myl7:mCherry.

For replacement of the myl7:mCherry secondary reporter with he1.1:YFP, the elavl3:hArr-TEV sequence were amplified from the elavl3:hArr-TEV; myl7:mCherry plasmid using primers 183 and 184. The he1.1:YFP sequence was amplified from p3E_he1a:eYFP (Addgene #113879) using primers 185 and 186. Both PCR products were assembled with the Tol2 Gateway plasmid backbone (pDestTol2pA2-U6:gRNA; Addgene #63157) using Hi-Fi DNA Assembly to obtain elavl3:hArr-TEV;he1.1:YFP.

hArr-TEV was also placed under the control of Xla.Tubb2b by amplification of this sequence from elavl3:hArr-TEV;myl7:mCherry using primers 203 and 51, with flanking attB1 and attB2 sequences, respectively. A Gateway recombination reaction was performed using BP Clonase II with the PCR product and donor vector pDONR221 (Invitrogen, #12536017) to obtain pME_hArr-TEV. LR Clonase II was used to recombine p5E_nbt, pME_hArr-TEV and p3E_he1a:eYFP (Addgene #113879) into the Tol2 Gateway Destination Vector (^69^) (#394) to obtain Xla.Tubb2b:hArr-TEV;he1.1:YFP.

To generate the QUAS:H2B-GCaMP6s; he1.1:CFP plasmid, an H2B-GCaMP6s middle entry clone was prepapred from Addgene plasmid #59530 and subsequently recombined with p5E_QUAS (Addgene, #61374) and p3E_he1a:ECFP (Addgene, #113880) into a Tol2 Destination Vector (^69^) (#394).

### Microinjection

Progeny from matings between Gal4 driver lines and the *Tg(QUAS:mApple-CAAX; he1.1:mCherry)^c6^*^36^ reporter line were microinjected at the 1-cell stage with plasmids for all *trans-*Tango components (10xUAS-E1B:sGCG-ICAM-(1235)Nrxn1b; cryaa:mCherry, *elavl3-hArr-TEV; he1.1:eYFP, elavl3-hGCGR-TEVcs-QF; he1.1:CFP)* together with Tol2 transposase mRNA (25 ng/μl). Tol2 transposase mRNA was produced by digesting pCS-zT2TP (^71^) with *Notl* and using the mMESSAGE mMACHINE SP6 Transcription kit (AM1340, Thermo Fisher Scientific) followed by phenol/chloroform-isoamyl extraction. Ligand and arrestin constructs (25 ng/µl) and the receptor construct (10 ng/ µl or 25 ng/µl or 125 ng/µl) were pooled in a 5 µl injection mixture with water and phenol red (.2%). At 24 hpf, injected embryos were transferred to 0.003% PTU and system water and screened under an Olympus MVX10 Macro Zoom fluorescence microscope for labeling of hatching gland cells. Embryos positive for YFP, CFP, and mCherry labeled hatching gland cells were reared to 4 dpf, then screened for mCherry labeling of the lens and the presence of the UAS:GFP reporter. At 6 dpf, larvae were screened on the Olympus microscope for mApple-CAAX labeling in the CNS. Those that were positive were individually mounted in 1% low melting point agarose (50100, Lonza) and imaged on a Zeiss LSM 980 confocal microscope using a 20x (NA=0.5) water immersion lens.

Whole mount in situ hybridization

Embryos from transgenic lines were presorted for labeling of appropriate secondary markers (refer to Fig. 1b) and, at 6 dpf, larvae were fixed in 4% paraformaldehyde (PFA; Sigma-Aldrich) in 1X phosphate-buffered saline (PBS) at 4°C overnight and then dehydrated with methanol. To generate probes, primers amplifying a 200-500 bp stretch of DNA sequence corresponding to each *trans*-Tango component were designed using Primer3 software (https://primer3.org/). The location, size of DNA template, and primer sequences for the ligand, receptor and Arrestin transgenes are provided in Table 2. The *trans*-Tango plasmids were used as templates to generate PCR products, which were purified using a PCR cleanup kit (QIAGEN QIAquick PCR purification kit #28104). Reverse primers included a promoter for in vitro transcription using SP6 RNA polymerase (Roche RPOLSP6RO) with incorporation of UTP-digoxigenin (Roche 11093274910). RNA was purified using G-50 columns (Cytiva 27533001) and diluted to 30 ng/ml. Larvae were rehydrated in PBS, permeabilized using Proteinase K (Roche 03115828001), post fixed in 4% PFA, and hybridized in 50% formamide containing 30 ng of the RNA probe at 70°C. Probes were detected using alkaline phosphate-conjugated digoxigenin antibody (Sigma Aldrich 11093274910) and visualized after incubation with substrate solution containing NBT (4-nitro blue tetrazolium, Roche 11383213001) and BCIP (5-bromo-4-chloro-3-indolyl-phosphate, Roche 11383221001). Larvae were mounted in 100% glycerol under coverslips and bright-field images captured using a Leica DFC500 camera mounted on a Zeiss Axioskop

**Table 2:**
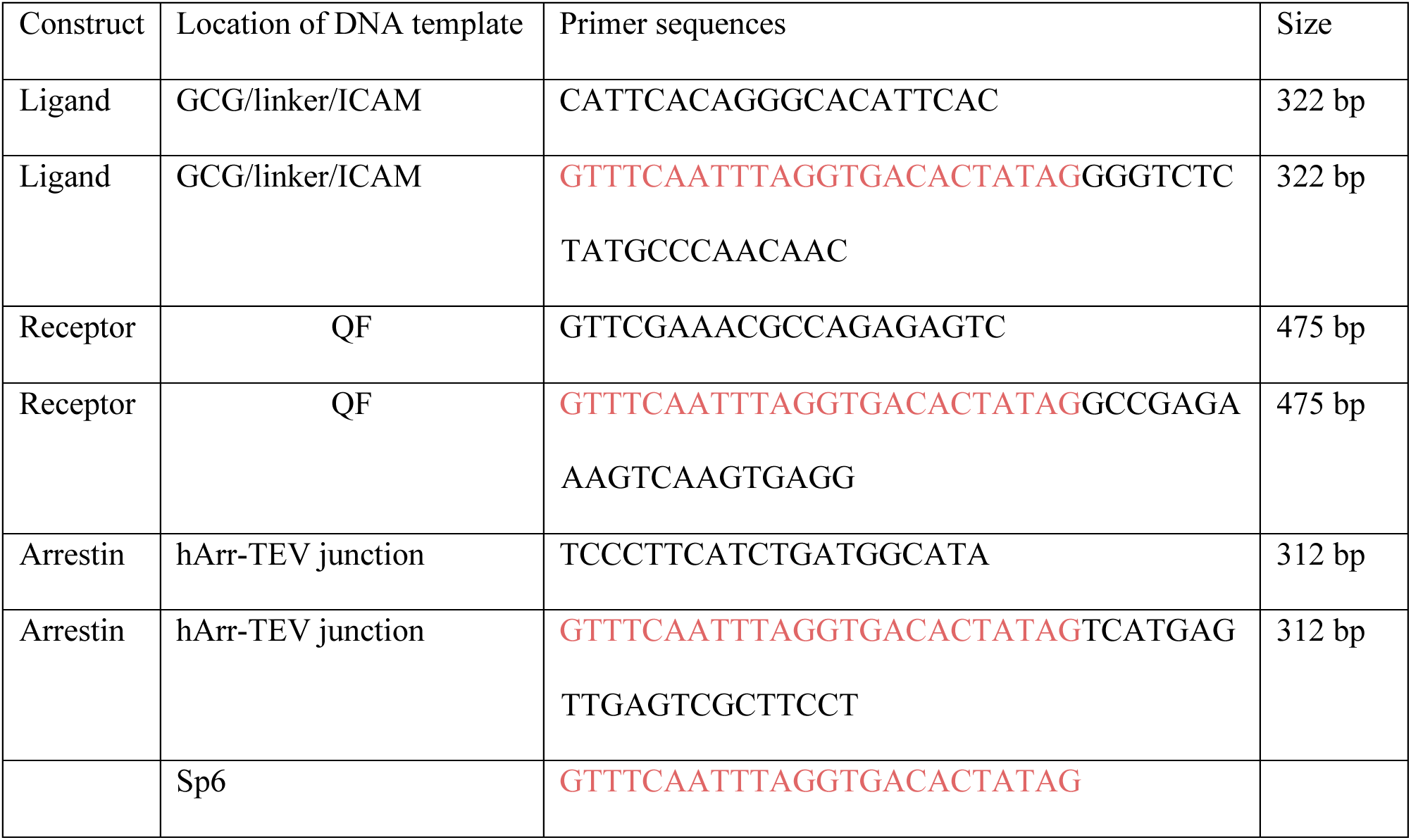
Primers for synthesis of probes for RNA in situ hybridization

### Immunofluorescence

Larvae were raised to 6 dpf in 0.003% PTU in system water, anesthetized using tricaine (0.02%, Sigma Aldrich E10521), and fixed in 4% paraformaldehyde overnight at 4°C. They were cryoprotected at 4°C overnight, first in 10% sucrose and then in 30% sucrose. Specimens were individually mounted nosedown in optimal cutting temperature (OCT) compound in plastic molds and frozen at -80°C. Sections (25 µm) were prepared on a HM525 NX Cryostat (Epredia), layered onto Superfrost Plus slides (Fisher Scientific), and dried overnight in the dark at room temperature. Subsequently, sections were post-fixed in 1% paraformaldehyde, washed 3 times each in PBS and PBST, and incubated in blocking solution containing PBST and 1% bovine serum albumin at room temperature for 1 hour. The sections were then incubated overnight at 4°C in primary antibody diluted in blocking solution with 5% normal goat serum (Abcam, ab7481). Antibodies were directed against GFP (Abcam, ab13970, 1:500), HuC/D (Thermo Fisher Scientific, A-21271, 1:40), PKCa (Santa Cruz Biotechnologies, sc-17769, 1:100) and zpr-3 (ZIRC, 1:100). Following three washes in PBST, sections were incubated at room temperature for 1 hour with either goat anti-mouse Alexa Fluor 647 (Thermo Fisher Scientific, a32728) or goat anti-chicken Alexa Fluor 488 (Thermo Fisher Scientific, a11039) secondary antibody diluted 1:1000 in PBST: Slides were washed in PBST, incubated 10 minutes with DAPI stain (1:1000 in PBS, Sigma Aldrich D9542-10MG), and washed in PBS before incubation in 10% glycerol overnight at 4°C. Sections were covered in 30% glycerol for at least one hour at room temperature before mounting in 40% glycerol.

### Confocal imaging

Larvae were anesthetized with 0.02% ethyl 3-aminobenzoate methanesulfonate salt (Tricane, (E10521, Ethyl 3-aminobenzoate methanesulfonate) and mounted in 1% low melting point agarose (50100, Lonza) in a 60 mm x 15 mm petri dish. Once solidified, system water containing PTU (0.003%) and Tricaine (0.02 %) was added to cover the agarose. Larvae were imaged using a Zeiss LSM 980 with Airyscan 2 under a 20X (NA 0.8) water immersion objective. Images were taken in Z-stacks and processed using Fiji software. For tracing the morphology of individual neurons labeled by *trans*Tango transsynaptic signaling, 3D renderings were also performed using Fiji (^72^). Image files were extracted using *File -> Split channels -> image -> adjust -> threshold -> invert -> save as -> Tiff*.

### Image registration

Confocal images of zebrafish larval brains were captured in two channels, one for the reference brain (GFP) and one for *trans*-Tango labeling (mApple). The reference larva was select based on orientation, straightness of head and lack of tilting and all other datasets (larvae) were registered to the reference channel (ptf1a/islet2b) of the reference larva using ANTs (Advanced Normalization Tools). We applied the code from Marquart et al. (^73^) with minor modifications to streamline the computation.

Computation was performed using the high-performance computer cluster managed by Research Computing at Dartmouth College.

~~~
antsRegistration --dimensionality 3 --verbose 1 --float 1 -o
[${file//01.nrrd/01_},${file//01.nrrd/01_warped.nrrd}] --use-histogram-matching 0 –interpolation
WelchWindowedSinc --initial-moving-transform [${refbrain//01.nrrd/01.nrrd},${file//01.nrrd/01.nrrd},1]
-t rigid[0.1] --metric MI[${refbrain//01.nrrd/01.nrrd},${file//01.nrrd/01.nrrd},1,32,Regular,0.25] -c
[200x200x200x0,1e-8,10] --shrink-factors 12x8x4x2 -s 4x3x2x1vox -t Affine[0.1] –metric
MI[${refbrain//01.nrrd/01.nrrd},${file//01.nrrd/01.nrrd},1,32,Regular,0.25] --convergence
[200x200x200x200x10,1e-7,10] --shrink-factors 12x8x4x2x1 -s 4x3x2x1x0vox;
antsApplyTransforms --dimensionality 3 --float 1 -v 0 -n WelchWindowedSinc -i ${file//01.nrrd/02.nrrd}
-r ${refbrain//01.nrrd/02.nrrd} -o ${file//01.nrrd/02_warped.nrrd} -t ${file//01.nrrd/01_1Warp.nii.gz} -t
${file//01.nrrd/01_0GenericAffine.mat};
~~~

To compare *trans-*Tango labeling between different datasets, we devised an algorithm to assign a unique color to the neurons of individual larvae by dividing the spectra from violet to red (0-360 degrees) with the number of individuals (n) in the Hue, Saturation, Lightness (HSV) color system. The registered CNS labeling for each larva was then plotted in a single image stack.

### Optogenetics and calcium imaging

Embryos from *Tg(isl2b:Gal4-VP16); Tg(UAS:ReaChR-Tag-RFPT)^jf^*^50^ intercrosses were injected with the QUAS:Hsa.H2B-GCaMP6s; he1.1:CFP plasmid along with the *trans*Tango ligand, receptor, arrestin plasmids at the 1-cell stage. Larvae containing all secondary markers were allowed to develop to 6 dpf, at which time they were paralyzed by a 1 min immersion in α-bungarotoxin (20 µl of 1 mg/ml solution in system water, B1601, ThermoFisher Scientific) and flushed with fresh system water. Individual larvae were mounted in a droplet of 1% low melting agarose (50100, Lonza) in the middle of a 60 mm x 15 mm Petri dish and immersed in system water after the agar solidified. For calcium imaging experiments, the xyt acquisition mode was used to capture images using a 20X (NA=0.5) water image objective on a Zeiss LSM 980. Images containing the optic tectum in the field of view were acquired with the 488 nm laser at 310 x 310-pixel resolution and a rate of 2.6 Hz while stimulating the left or right retina with 561 nm light.

Calcium transients were recorded in a series of 200 frames prior to retinal activation, 20 frames during activation with the 561 nm laser, and 150 frames following activation.

Image files were extracted in Fiji using *File -> Save as -> Image Sequence Files*. Frames were imported to MATLAB (Mathworks Inc.), and mean fluorescence intensities for individual regions of interest (ROI; areas adjacent to the left and right optic tectum) were calculated. For each larva, a highcontrast image was generated by calculating a maximum intensity projection for the series of images generated in Fiji. ROIs were drawn manually using the MATLAB *roipoly* function and the mean fluorescence intensity of pixels within each ROI was calculated. The change in fluorescence (ΔF/F) was calculated according to the following formula

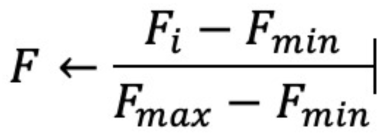

where Fi represents the mean fluorescence intensity in an ROI at each time point, and Fmax and Fmin are the maximum and minimum fluorescence values, respectively, for that ROI during the recording period. To calculate total activity for each larva before and after activation, the ΔF/F was averaged across the ROIs of each larva and total activity was obtained for a period by calculating the area under the curve using the MATLAB *trapz* function.

## Acknowledgements

We thank Misha Ahrens, Tammy Kaminy, Kristen Kwan, Tim Mulligan and Jeff Mumm for sharing reagents and transgenic lines. We are grateful to Eric Horstick for advice on codon optimization and to Emma Spikol and Ahmed Abdelfattah for expert guidance on calcium imaging experiments. Special thanks are extended to Pat Robison for microscopy support and Bethany Malskis and Jaden Devine Brilliant for animal care. This work was supported by a Hanna H. Gray fellowship to CEC and RF1MH123213 from the NIH Brain Initiative to MEH, GB, JL and DR.

