## Supplemental Figures for "Transsynaptic labeling and transcriptional control of zebrafish neural circuits"

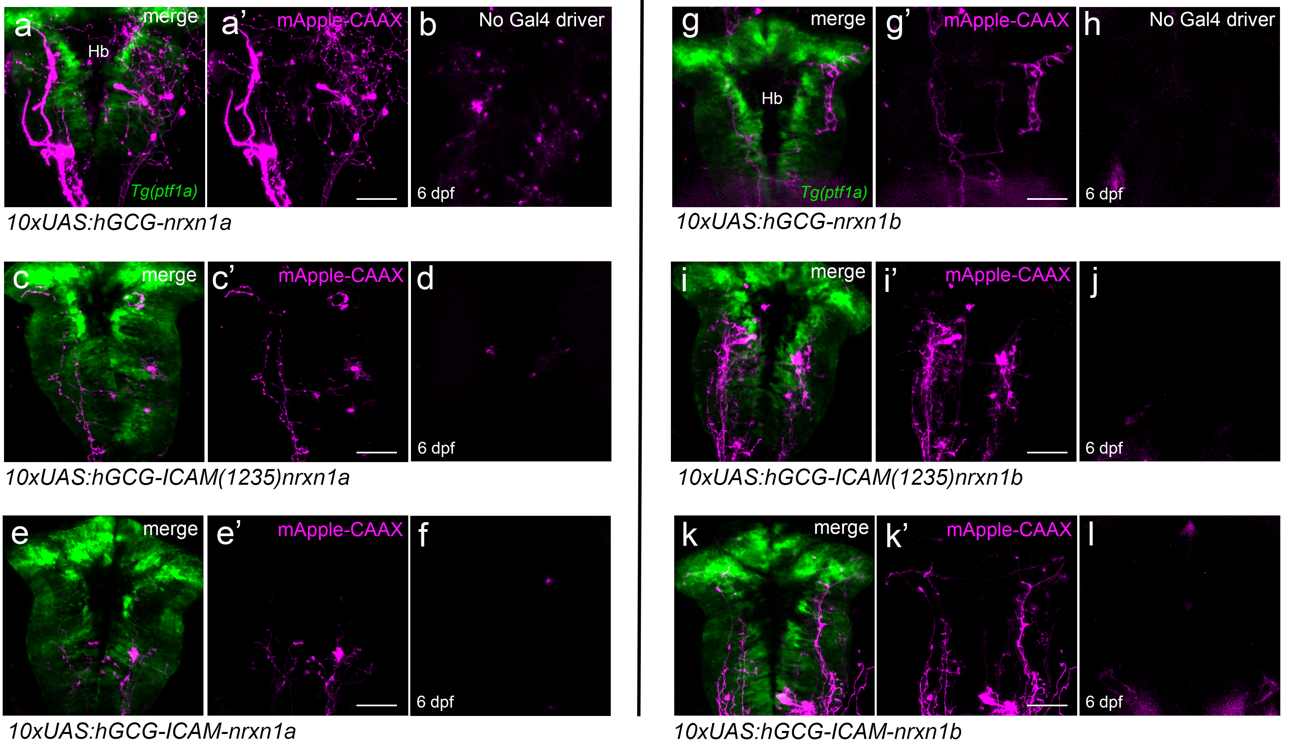

Supplemental Fig. 1. Optimization of the *trans-*Tango ligand

Six different ligand constructs were injected along with plasmids encoding the *trans-*Tango receptor, arrestin-TEV and Tol2 transposase RNA into 1-cell zebrafish embryos, the progeny of *Tg(QUAS:mApple‑CAAX)* and the *Tg(ptf1a:Gal4‑VP16)* driver line that promotes strong expression of *Tg(UAS:GFP)* in the hindbrain. (a-f) Ligands containing the transmembrane domain of zebrafish *nrxn1a* and varying lengths of mouse ICAM. (a-a’) *10xUAS-E1B:sGCG-nrxn1a* produced consistent *trans*-Tango labeling in the hindbrain (n= 20/25) but also (b) nonspecific labeling (n=15/25). (c-c’) *10xUAS-E1B:sGCG-ICAM(1235)-nrxn1a* produced consistent hindbrain labeling (n= 17/25) and (d) nonspecific labeling (n=11/25). (e,e’) *10xUAS-E1B:sGCG-ICAM-nrxn1a* produced fewer *trans*-Tango labeled larvae (n= 6/25) and (f) nonspecific labeling (n=5/25). (g-l) Ligands containing the transmembrane domain of zebrafish *nrxn1b* and varying lengths of mouse ICAM. (g-g’) *10xUAS-E1B:sGCG-nrxn1b* produced consistent *trans*-Tango labeling (n= 15/25) and (h) substantial nonspecific labeling (n= 12/25). (i,i’) *10xUAS-E1B:sGCG-ICAM(1235)-nrxn1b* produced consistent labeling in the hindbrain (n= 23/25) and (j) minimal nonspecific labeling (n=2/25). (k,k’) *10xUAS-E1B:sGCG-ICAM-nrxn1b* produced *trans*‑Tango labeling (n= 21/25) and (l) nonspecific labeling (n=14/25). Scale bars, 50 μm.

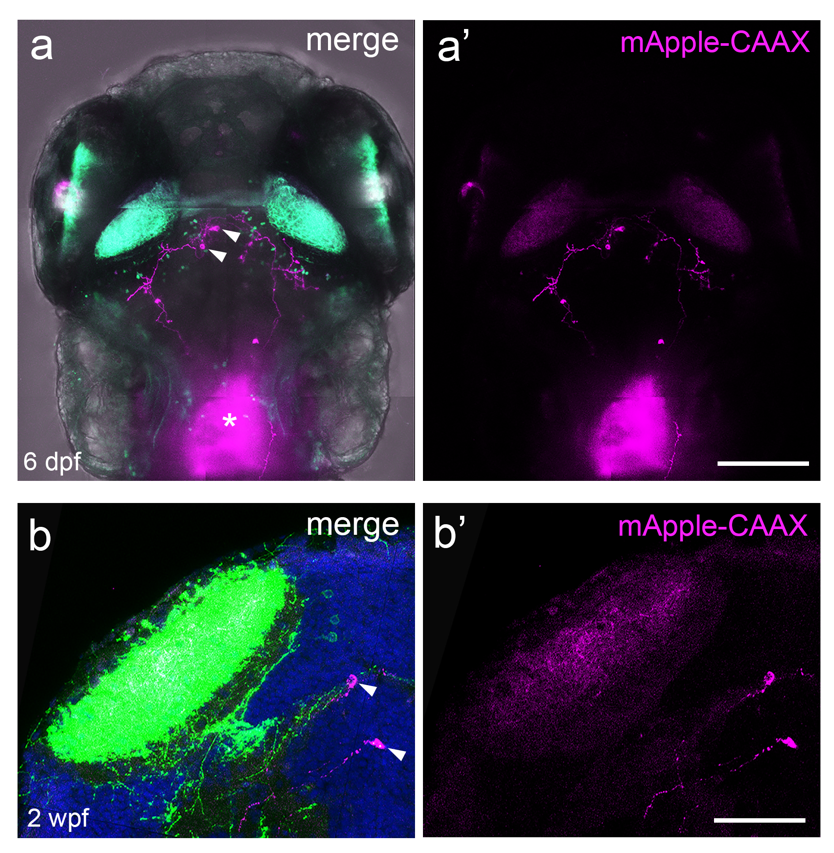

### Supplemental Fig. 2. Monitoring synaptic connections over time

(a, a’) Dorsal view of mApple-CAAX labeled neurons in the optic tectum (arrows) of a 6 dpf larva bearing *Tg(isl2b.2:Gal4; myl7:tagRFP); Tg(UAS:GFP)*, *Tg(QUAS:mApple-CAAX;he1.1:mCherry)* and all *trans*-Tango components. Diffuse red fluorescent protein labeling (asterisk) is due to a secondary marker (*my17:RFP*) expressed in the heart. (b,b’) In the same animal, labeling of the tectal neurons indicated in (a; arrows) persisted at 2 weeks post-fertilization (wpf). Scale bars, 50 μm in a,a’ and 100 μm in b,b’.

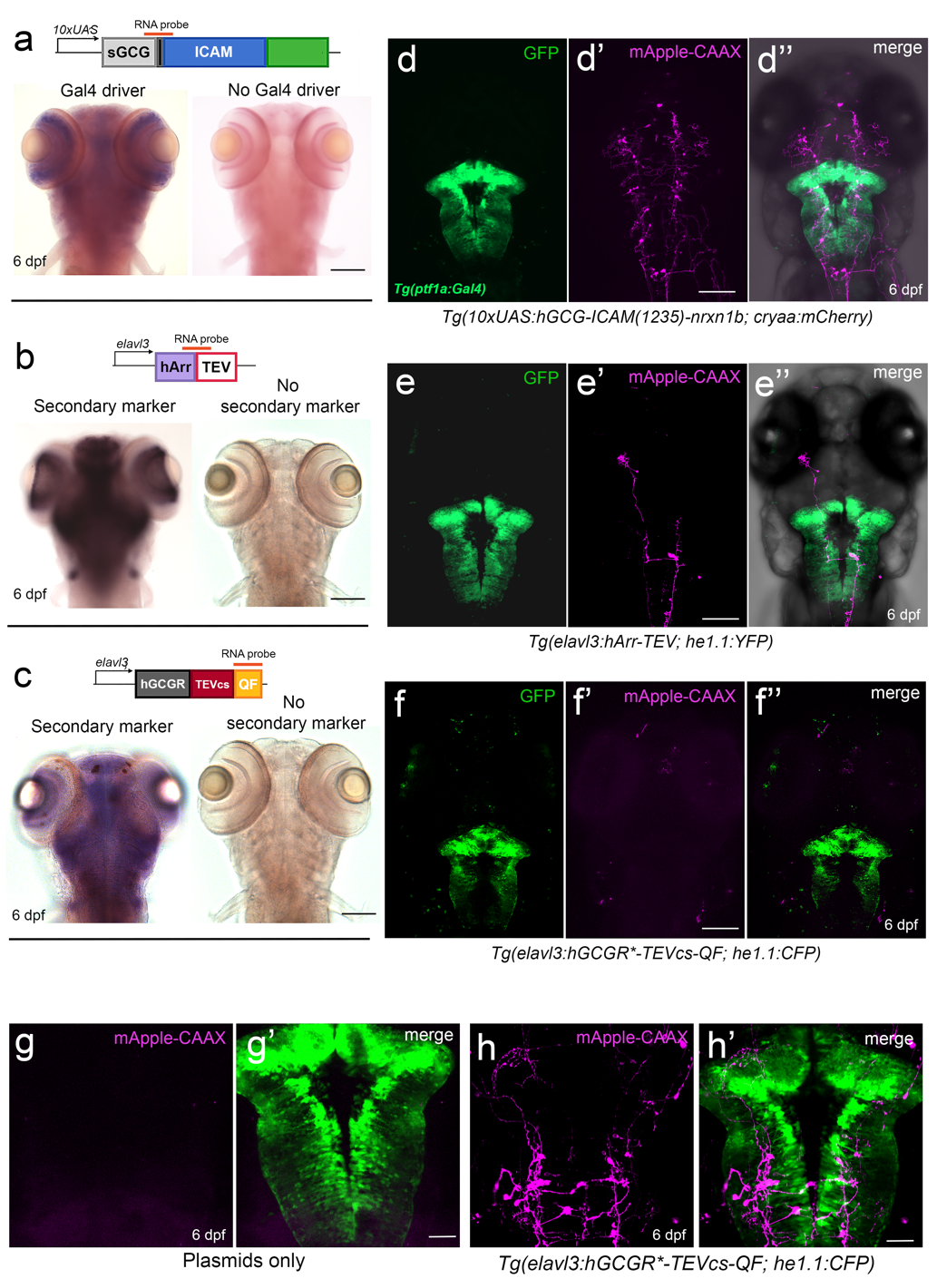

### Supplemental Fig. 3. Generation of *trans-*Tango transgenic zebrafish

Heterozygous fish bearing transgenes for *trans*-Tango components and Tg*(QUAS:mApple-CAAX; he1.1:mCherry)* were mated with the (*TgBAC(pt1fa-Gal4-VP16; UAS:GFP)^jh16^* driver line and resultant embryos injected with plasmids for the other *trans*-Tango reagents and Tol2 RNA. (a) RNA *in situ* hybridization indicates that the expression of UAS:sGCG-ICAM(1235)-Nrxn1b resembles the transcription pattern of the endogenous *ptf1a* gene (Kani et al., 2010). Both the (b) *Tg(elavl3:hArr-TEV; he1.1:YFP)^cd30^* and (c) *Tg(elavl:hGCGR-TEVcs-QF; he1.1:CFP*) stable lines are expressed broadly throughout the CNS. (d,d’’) *Tg(10xUAS-E1B:sGCG-ICAM(1235)nrxn1b; cryaa:mcherry)* larvae showed *trans*-Tango labeling in the hindbrain (n=257/312) consistent with that observed when the ligand is introduced by plasmid injection. (e,e’’) *Tg(elavl3:hArr-TEV; he1.1:YFP)* larvae had similar *trans*-Tango labeling in the hindbrain (n= 132/164). (f,f’’) mApple-CAAX labeling was not observed in the hindbrain (n= 0/276) with the receptor line *Tg(elavl:hGCGR-TEVcs-QF; he1.1:CFP*) or when (g,g’) receptor plasmid was injected at a sub-threshold concentration (10 ng/μl) along with the ligand and arrestin-TEV plasmids (25 ng/µl) (n=25). (h,h’) However, the same concentration of receptor plasmid was effective at producing extensive mApple-CAAX labeling in the hindbrain (n= 21/25) when injected into embryos bearing *Tg(elavl:hGCGR-TEVcs-QF;he1.1:CFP)* along with the ligand and arrestin-TEV plasmids (25 ng/µl). Scale bars, 20 µm and 50 μm.

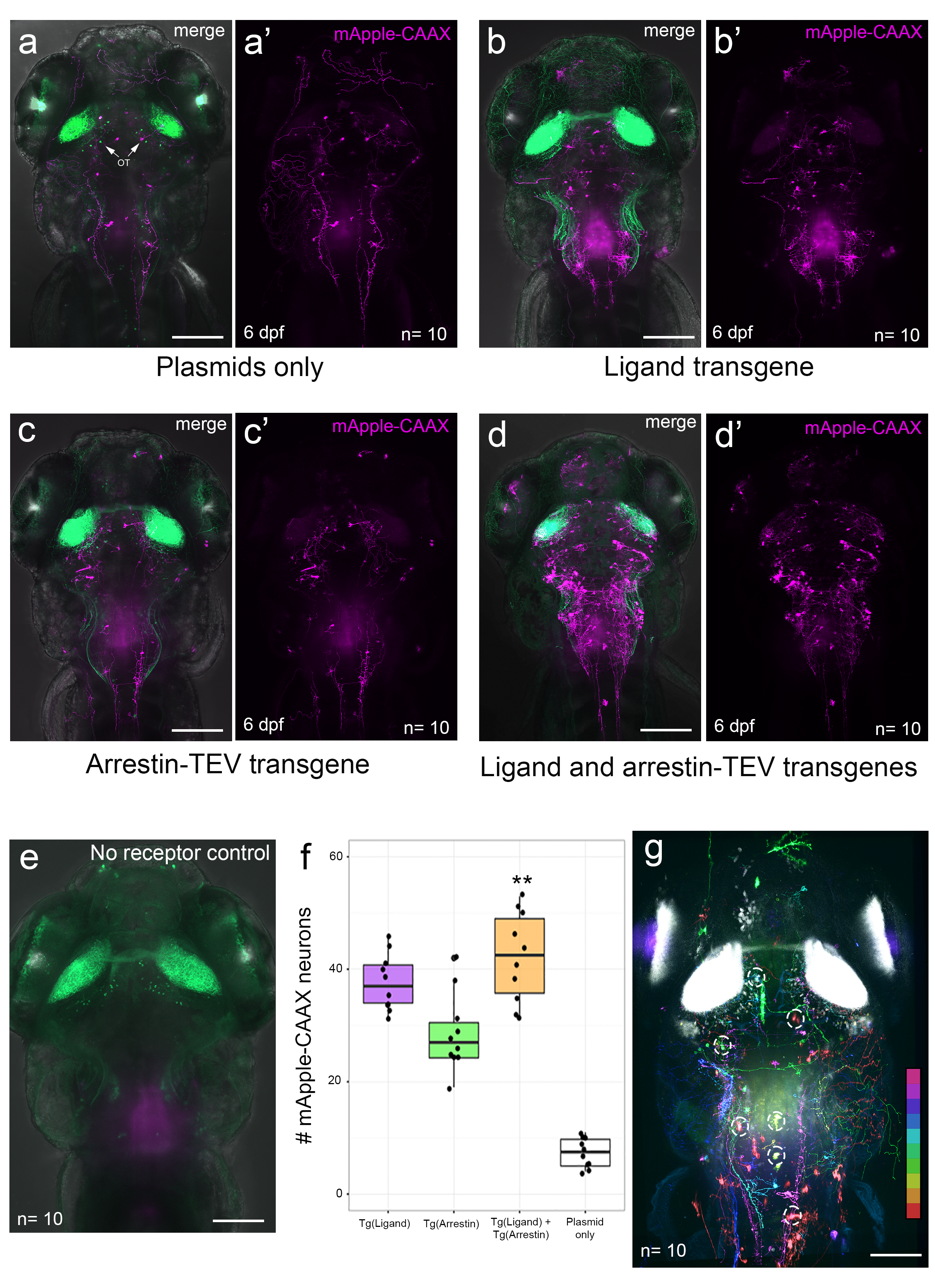

Supplemental Fig.4. *trans*-Tango labeling in stable transgenic lines.

Robust mApple-CAAX labeling in *Tg(isl2b.2:Gal4; myl7:tagRFP*) 6 dpf larvae bearing both the ligand and arrestin-TEV stable transgenes and the *Tg(UAS:GFP)* and *Tg(QUAS:mApple-CAAX)* reporter lines. (a,a’) Control larvae from injections of plasmids for all *trans*-Tango components (n = 10). (b,b’) *Tg(10xUAS-sGCG-ICAM(1235)-nrxn1b; cryaa:mcherry)* larvae injected with the receptor and arrestin-TEV constructs (n = 10). (c,c’) *Tg(elavl3:hArr:TEV* ; he1.1:YFP) larvae that had been injected with the receptor and ligand constructs (n = 10). (d,d’) Larvae transgenic for both the ligand and arrestin-TEV transgenes that had been injected with the *trans-*Tango receptor construct (n = 10). (e) Uninjected controls (n = 10). (f) On average, 45 neurons were labeled with mApple-CAAX in larvae that bore both the ligand and arrestin-TEV stable transgenes compared to those from injections of all *trans*-Tango components (on average, 11 neurons per sample). (g) Image registration of mApple-CAAX labeled neurons in the optic tectum and hindbrain of ten *Tg(isl2b.2:Gal4;myl7:tagRFP;UAS:GFP)*; *Tg(10xUAS-sGCG-ICAM(1235)-nrxn1b; cryaa:mcherry*); *Tg(elavl3:hArr:TEV* ; he1.1:YFP) larvae injected with the receptor construct. Patterns of mApple-CAAX labeled neurons are pseudocolored differently for each larva and dashed circles indicate some cells that were labeled in multiple samples. Scale bars, 20 μm.
